# Plastic-elastomer heterostructure for robust flexible brain-computer interfaces

**DOI:** 10.1101/2025.04.29.651325

**Authors:** Xinyi Lin, Xinhe Zhang, Zheliang Wang, Juntao Chen, Jaeyong Lee, Ariel J. Lee, Hang Yang, Antoine Remy, Hao Shen, Yichun He, Hao Zhao, Xuyue Zhang, Wenbo Wang, Almir Aljović, Joost J. Vlassak, Nanshu Lu, Jia Liu

## Abstract

Electronics for neural signal recording must be robust across multiple and deep brain regions while preserving tissue-level flexibility to ensure stable tracking over months or years. However, existing electronics cannot simultaneously achieve robustness and tissue-level flexibility, limiting their potential for customizable and scalable neuroscience research and clinical applications. Here, we introduce FlexiSoft, an electronic platform based on a plastic-elastomer heterostructure that uniquely integrates mechanical robustness and tissue-level flexibility. Compared to conventional flexible electronics of similar thickness, the FlexiSoft platform demonstrates an order-of-magnitude improvement in both mechanical robustness (critical energy release rate) and flexibility (flexural rigidity). Leveraging these mechanical advantages, we developed FlexiSoft probe for robust implantation, demonstrated by its ability to withstand repeated insertion and removal, as well as to reach centimeter-scale depths comparable to those in the human brain. The platform enables long-term recording from the same neurons across the hippocampus (HPC) and primary motor cortex (M1) during a months-long motor learning task, thereby revealing long-term dynamic changes in neuronal firing patterns. Additionally, FlexiSoft’s unique robustness and flexibility enable curved implantation routes, opening new directions of customizable implantation pathways. In summary, we present FlexiSoft as a novel, robust, and tissue-level flexible heterostructure electronics platform that advances flexible brain-computer interfaces (BCIs) with strong translational potential for neuroscience and clinical applications.

## Main text

Tracking activity from the same neurons across multiple deep brain regions at single-cell resolution over extended periods is critical for understanding learning and memory^1–4^, disease progression^5^, and therapeutic development^6^. However, current neural recording platforms face significant limitations. High-density rigid silicon-based electrode arrays^7–9^ frequently exhibit probe drift^10^ and induce chronic tissue damage^11^, owing to their mechanical mismatch with soft brain tissue^12^. Attempts to reduce tissue damage by thinning silicon substrates compromise the mechanical robustness of these devices, hindering reliable implantation into deep brain regions^13,14^. As a result, thicker probes are often required, which further exacerbates the mechanical mismatch and associated challenges^15^.

To overcome these limitations, flexible electronics have been developed using ultrathin polymer substrates^16–19^, achieving tissue-level flexibility that minimizes immune responses and probe drift. However, these ultrathin structures are mechanically fragile, rendering them difficult to handle^20^ and prone to cracking during fabrication, transfer, and implantation^21,22^. This fragility limits implantation yield and complicates reliable insertion into multiple deep brain regions. Moreover, the brittleness of ultrathin devices increases the risk of fractures during removal, potentially leaving fragments in brain tissue —a critical concern for clinical applications. To address these issues, soft electronics using elastomers as dielectric materials^11,17^ have been introduced, offering enhanced flexibility and reduced brittleness. However, the extreme stretchability of these elastomer-based devices often exceeds the mechanical limits of embedded metal interconnects and electrodes, leading to interconnect breakage during implantation, particularly when excessive stretching occurs^23^. Consequently, the fragility and handling challenges of both flexible and soft electronics remain major barriers to their widespread adoption.

An ideal brain probe would combine a soft yet non-stretchable substrate and dielectric layers with chronic stability in biofluidic environments. To address this need, we introduce the FlexiSoft neural probe—a robust, tissue-level flexible platform (Fig. 1a) that integrates soft elastomer layers for mechanical support with thin-film flexible layers to constrain stretchability. This plastic- elastomer heterostructure achieves a tenfold enhancement in both mechanical robustness, as quantified by critical energy release rate, and flexibility, as determined by flexural rigidity, while retaining a total thickness comparable to that of conventional flexible electronics. The soft, stretchable elastomer layer mitigates the formation and propagation of micro-cracks and, critically, suppresses channel cracking in the fragile plastic and thin metal layers —a primary failure mode in traditional flexible neural probes—while the flexible dielectric layers support and protect the thin metal interconnects and electrodes. Leveraging these unique mechanical properties, the FlexiSoft neural probe enables high-yield, repeated implantation and retrieval without mechanical failure. It also supports robust implantation across multiple brain regions (Fig. 1b). Importantly, by preserving tissue-level mechanical properties, FlexiSoft probes enable, for the first time, stable tracking of single-unit activity from the same neurons across multiple brain regions, including hippocampus (HPC) and primary motor cortex (M1), throughout a months-long learning task (Fig. 1c-d). Furthermore, the FlexiSoft neural probe supports customizable curved-path implantation for diverse neural recordings and enables centimeter-scale implantation depth consistent with human brain dimensions, demonstrating strong potential for clinical translation.

**Fig. 1.**
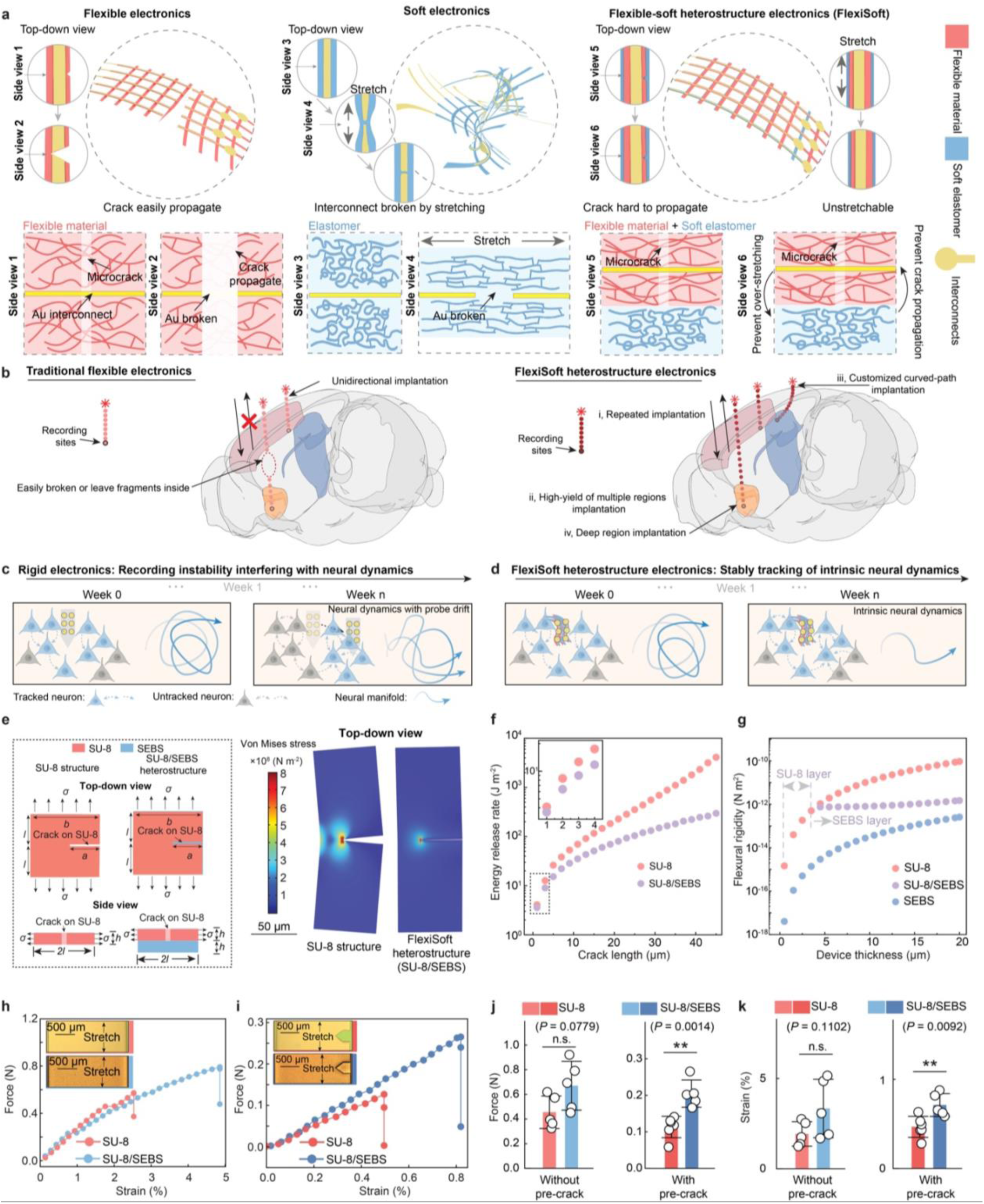
FlexiSoft platform: A robust flexible-soft heterostructure electronics platform for stable brain-computer interfaces. a, Comparative schematics of conventional thin-film flexible, soft, and FlexiSoft devices. FlexiSoft device integrates a soft elastomer layer with a flexible polymer, enhancing mechanical support, minimizing crack propagation, and preventing structural failure while preserving tissue-level flexibility. **b,** Key features of FlexiSoft device include improved resistance to breakage, high-yield implantation across deep and multiple brain regions, and support for customizable curved implantation pathway, enabling long-term stable tracking activity from the same neurons. **c-d,** Schematic illustrations of the importance of stable neural recordings. (**c**) Rigid electronics cause probe drift, mixing intrinsic neural dynamics with recording instability. (**d**) FlexiSoft device enables stable tracking activity from the same neurons, isolating intrinsic long-term neural dynamics. **e,** Finite element analysis (FEA) of a thin-film, sheet-like neural probe shank. Left: Schematics showing the top-down and side views of a SU-8 structure and a SU-8/SEBS heterostructure, both with a crack on the SU-8 layer (2*l* = 200 μm, *h* = 4 μm, *b* = 70 μm, *σ* = 50 MPa). Right: von Mises stress distribution around a 45 μm crack in a structure of 70 μm in width and 200 μm in length, composed of either SU-8 alone (4 μm thick) or a SU- 8/H-SEBS heterostructure (4 μm SU-8 + 4 μm SEBS) under identical tensile load (constant far-field uniaxial stress). **f,** Energy release rate as a function of crack length for SU-8 and SU-8/H- SEBS structure under the same geometry and loading conditions as in (**e**). The inset demonstrates that the FlexiSoft structure exhibits a significantly lower energy release rate across crack length, even when microcrack dimensions are comparable to SU-8 thicknesses. **g,** Flexural rigidity as a function of thickness for SU-8, H-SEBS and FlexiSoft structure. **h-i,** Representative tensile tests on intact (**h**) and pre-cracked (**i**) samples comparing SU-8 and FlexiSoft structure. **j-k,** Quantification of force at break (**j**) and strain at break (**k**) of SU-8 versus FlexiSoft structure under intact and pre-cracked conditions. Data are represented as mean ± SD, two-tailed t-test: \*\**P* < 0.01 (n = 6 sample ftests for SU-8 structure with pre-crack, n = 5 sample tests for others).

### Design of the FlexiSoft neural probe for enhanced robustness and flexibility

The core design principle of the FlexiSoft neural probe is that introducing an elastomer substrate into a thin-film probe effectively constrains crack opening and reduces the energy release rate (*G*) under the same loading, where *G* quantifies the energy available for crack propagation^24–27^. A lower *G* corresponds to greater resistance to crack growth. For a given material with a critical energy release rate (*Gc*), crack propagation occurs only when *G ≥ Gc*. Therefore, reducing *G* makes it less likely to exceed *Gc*, enhancing fracture resistance under the same loading conditions.

To test this concept, we combined an SU-8 thin film with a hydrogenated styrene-ethylene-butylene-styrene (H-SEBS) elastomer substrate. The mechanical behavior of the SU-8-only structure and FlexiSoft (SU-8/H-SEBS) heterostructure were evaluated using finite element analysis (FEA, Methods). First, we analyzed the von Mises stress distribution in both structures under uniaxial tensile stress with varying crack lengths (Fig. 1e and Extended Data Fig. 1a-b). Under the same far-field tensile stress, the SU-8 structures exhibited a broader stress concentration zone around the crack compared to the SU-8/H-SEBS heterostructure. At a crack length of 45 μm, the simulated maximum von Mises stress in the SU-8 structure was 1.8 times greater than that in the SU-8/H-SEBS structure, indicating that the elastomer substrate in the FlexiSoft design effectively mitigates stress concentration. Then, to quantify their mechanical robustness, we calculated the energy release rate (*G*) under uniaxial load as a function of crack lengths using FEA (Fig. 1f). As crack length increased from 1 μm to 45 μm, the disparity in *G* between the two structures increased. At a 45 μm crack length, *G* in the SU-8 structure was 13.8 times higher than that in the SU-8/H-SEBS structure. These results demonstrate that adding an H-SEBS elastomer layer significantly enhances the robustness of the heterostructure by (1) reducing stress concentration around cracks and (2) decreasing *G* for crack propagation. All simulations and the theoretical solution yielded consistent results, indicating the simulation results are stable and accurately reflect the robustness of SU-8/H-SEBS heterostructure (Extended Data Fig. 1c, Supplementary Discussion 1 and Supplementary Fig. 1).

Next, we examined how adjusting the Young’s modulus and thickness of the H-SEBS substrate modulates *G* to inform device design. First, we examined *G* as a function of substrate Young’s modulus and SU-8 crack length under uniaxial tensile stress (Extended Data Fig. 1d-e). As Young’s modulus of the H-SEBS substrate increases, *G* decreases slowly for a given crack length. At a 45 μm crack length, *G* with a 10 MPa H-SEBS layer was 90.9% and 83.6% of those with 5 and 1 MPa H-SEBS layers, respectively, demonstrating that stiffer elastomer substrates slightly enhance structural robustness. Second, we varied H-SEBS thickness from 1 to 20 μm and found that thicker layers substantially reduced *G* (Extended Data Fig. 1f-g). For a 45 μm crack length, the *G* value of a 20 μm H-SEBS layer was only 38.3% of that in a 1 μm layer, showing an approximate threefold decrease in *G*. Together, these results indicate that increasing both the Young’s modulus and thickness of the H-SEBS substrate enhances the robustness of the FlexiSoft structure by minimizing *G*.

However, optimizing the FlexiSoft design requires balancing robustness with flexibility, characterized by flexural rigidity, *D*. While increasing the Young’s modulus and thickness of the H-SEBS layer reduces *G*, it also increases *D*, compromising flexibility. To maintain flexibility, we calculated *D* for SU-8, H-SEBS, and SU-8/H-SEBS heterostructures across thicknesses ranging from 1 to 20 μm (Fig. 1g, Supplementary Discussion 2-3). The *D* of the SU-8 structure increases rapidly with thickness, from ∼10^−12^ N·m^2^ at 4 μm to ∼10^−10^ N·m^2^ at 20 μm – a 125-fold increase. In contrast, the SU-8/H-SEBS heterostructure with a 4-μm-thick SU-8 layer can maintain *D* around ∼10^−12^ N·m^2^, when the total device thickness is increased from 5 to 20 μm.

Finally, we experimentally validated the enhanced robustness of the FlexiSoft structure through tensile stretching tests comparing SU-8 structures and SU-8/H-SEBS heterostructures, both with and without pre-cracks (Fig. 1h-k). During the stretching, cracks first appeared in the SU-8 layer, followed by the elongation of the SEBS layer and subsequent SU-8/SEBS delamination, indicating strong interlayer adhesion (Extended Data Fig. 2a-d). For structures without pre-cracks, the SU- 8/H-SEBS heterostructure exhibited approximately 1.5 times higher breaking force and 1.7 times higher breaking strain compared to SU-8 alone. When a pre-crack equivalent to 20% of the total width was introduced, SU-8/H-SEBS heterostructures still outperformed SU-8 structure, with breaking force and strain 1.8 and 1.5 times higher, respectively. These results confirm that the FlexiSoft heterostructure significantly enhances mechanical robustness compared to conventional thin-film flexible electronics.

### Fabrication of FlexiSoft neural probes

We developed a lithographic fabrication pipeline to integrate elastomer layers into a flexible neural probe (Methods). Conventional elastomers are easily swell in organic solvents and vulnerable to acid and base etching^28,29^, making them incompatible with multilayer microfabrication processes. To overcome these issues, we patterned the H-SEBS elastomer as the final fabrication step and deposited a 50 nm gold (Au) layer on top to enable precise patterning while protecting the elastomer from solvents, acids, and bases (Extended Data Fig. 3a). However, the large mismatch in thermal expansion coefficients between H-SEBS (149 ppm K^-1^) and Au (14.13 ppm K^-1^) led to cracking of the Au layer during thermal processing. To prevent this, we developed a low-temperature fabrication process in which all metal deposition, photoresist patterning, and reactive ion etching were performed below 90 ℃ (Methods).

The final FlexiSoft neural probe design incorporates (Fig. 2a): (i) Au interconnects encapsulated by top and bottom SU-8 layers; (ii) platinum (Pt)-coated electrodes for reduced interfacial impedance; (iii) guide holes for stereotaxic implantation; (iv) an H-SEBS support layer for enhanced fracture resistance. A representative FlexiSoft neural probe consists of five shanks, each containing six 20-μm-diameter electrodes spaced 10 μm apart, enabling simultaneous tracking of the same neurons across multiple electrodes (Fig. 2b). Contact profilometer measurements confirmed that the layer thicknesses matched the design: 2 μm for both the top and bottom SU-8 encapsulation layers and 4 μm for the H-SEBS layer (Extended Data Fig. 3b-d). Scanning electron microscopy (SEM) images and corresponding element mapping (Fig. 2c-f and Extended Data Fig. 4) confirmed the seamless integration of the H-SEBS and SU-8 layers. Imaging of FlexiSoft probes after releasing from the wafer confirmed their flexibility and structural integrity, with no delamination observed between the SU-8 and H-SEBS layers (Fig. 2g-h).

**Fig. 2.**
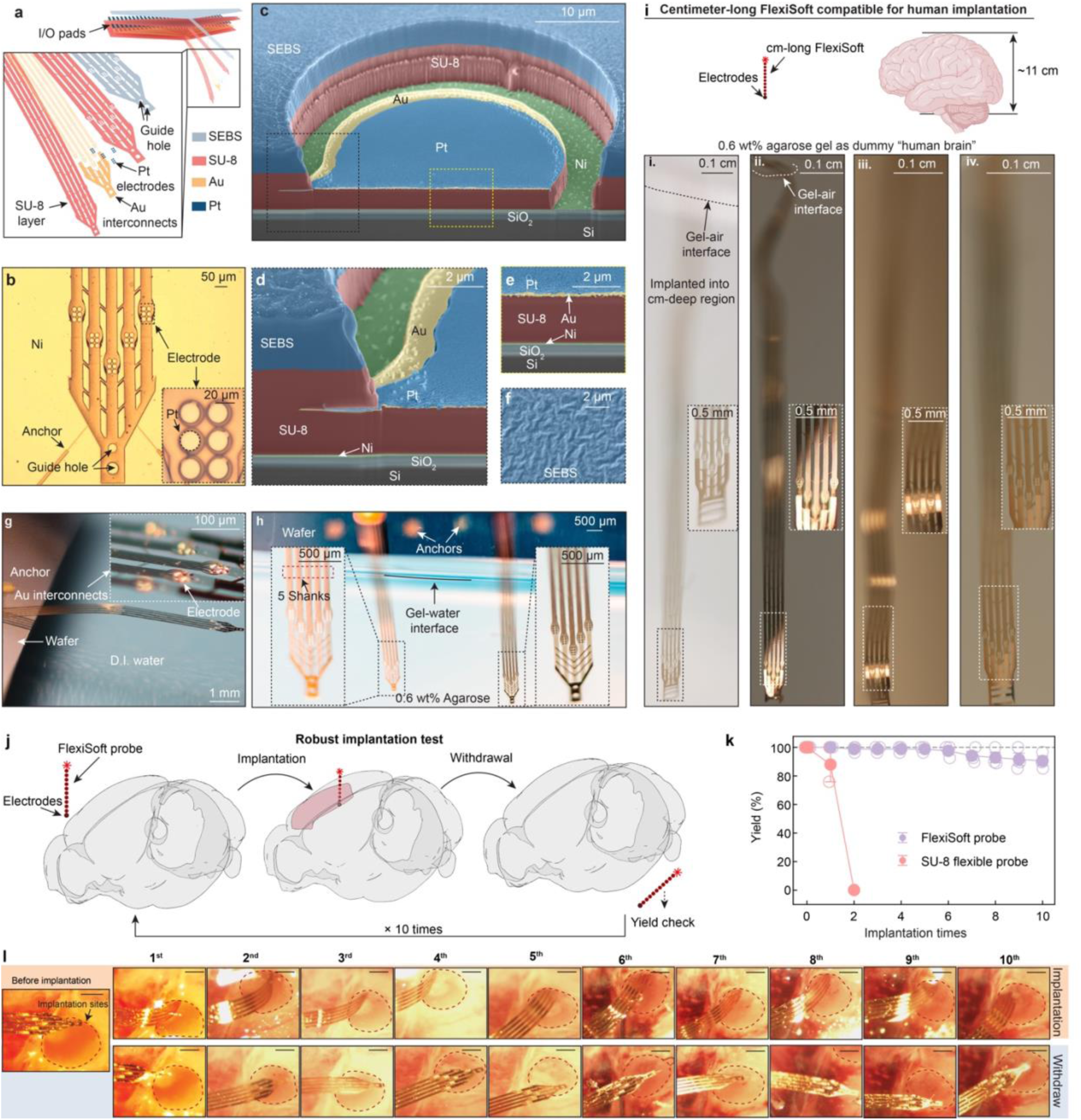
FlexiSoft neural probes for robust brain implantation. a, Schematic representation of the five-layer structure of a FlexiSoft neural probe. **b,** Bright-field (BF) microscopic image of an unreleased FlexiSoft probe on a Ni sacrificial layer. The zoomed-in view shows one shank of a six-electrode array. **c,** Scanning electron microscope (SEM) image showing an encapsulated electrode with pseudo-colored layers highlighting their material composition. **d**, Zoomed-in view of the black dashed box in (**c**) showing the interface between SU-8 and H-SEBS. **e**, Zoomed-in view of the yellow dashed box in (**c**) showing the multi-layered structure of a single electrode. **f**, SEM image showing the surface morphology of the H-SEBS soft elastomeric supporting layer. **g,** Photograph of a released FlexiSoft probe freely floating in deionized (D.I.) water. Insets: zoomed-in view of electrode arrays. **h,** Multi-probe, multi-region implantation in 0.6% agarose phantom gel, with a zoomed-in view of the implanted electrode arrays. **i,** Representative centimeter-long implantation of FlexiSoft in a “human brain” phantom gel (0.6% agarose gel) demonstrates its structural integrity after implantation at a scale relevant to the human brain. The zoomed-in view shows the intact flexible electrode arrays in the deep implanted region. **j,** Schematic representation of repeated implantation-retrieval cycles in a mouse brain used to assess the robustness of FlexiSoft probes. **k,** Dot plot showing the yield of electrodes over 10 repeated implantation cycles for three different FlexiSoft probes. Impedance values were measured after each implantation- retrieval cycle. **l,** BF images from a 10-cycle repeated implantation test, showing that the FlexiSoft probe remains structurally intact during 10 implantation-retrieval cycles. Scale bar: 500 μm.

To demonstrate the exceptional mechanical robustness of the FlexiSoft neural probe, we first implanted centimeter-long FlexiSoft probes into agarose gel, mimicking the size and mechanical stiffness of human brain tissue. Imaging results showed that the FlexiSoft probes maintained their structural integrity even after deep insertion comparable to the depth of human brain regions (Fig. 2i). Secondly, we conducted consecutive implantation and retrieval cycles in mouse brains, a critical requirement for clinical translation but one that conventional flexible probes typically fail to meet (Fig. 2j). Electrode impedance was measured after each implantation-retrieval cycle to assess electrode functionality^30,31^. Throughout ten consecutive cycles, FlexiSoft probes maintained stable performance, with channel yield remaining at 90.5 ± 3.3% after the tenth implantation (Fig. 2k, *n* = 3, mean ± SEM; Methods). In contrast, only one-third of conventional SU-8 flexible probes (*n* = 9) survived even the first implantation, and none remained intact after the second cycle. Brightfield (BF) images of a representative FlexiSoft probe after multiple implantation cycles confirmed an intact structure, with no device breakage inside the brain (Fig. 2l). Additionally, no delamination between SU-8 and H-SEBS layers was detected. These results demonstrate the exceptional mechanical robustness, reliability, and durability of FlexiSoft neural probes compared to conventional tissue-level flexible neural probes.

### Months-long stable tracking of single-unit action potentials from the same neurons across different brain regions

To evaluate whether the FlexiSoft probes allow stable tracking of single-cell activity over months-long implantation periods, we conducted both *in vitro* chronic impedance measurements and *in vivo* chronic recordings in mice. *In vitro* impedance measurements in saline over 20 weeks showed that electrode impedance values remained within the range of 1.3 ± 0.4 MΩ (Mean ± SD), consistent with previously reported values for implantable neural probes of similar electrode size and materials^16^ (Fig. 3a, Methods). Micro-CT imaging after 60 weeks of implantation in a mouse brain confirmed that FlexiSoft probes remained structurally intact in the mouse brain (Fig. 3b, Methods). For electrophysiological recordings, voltage signals were bandpass-filtered between 300-3000 Hz and referenced to a common reference electrode (Fig. 3c). Then, spike sorting was performed using MountainSort4^32^ on concatenated data from multiple sessions, followed by manual curation. In a representative mouse, we successfully tracked the same units across the electrode array over 126 days (Fig. 3d).

**Fig. 3.**
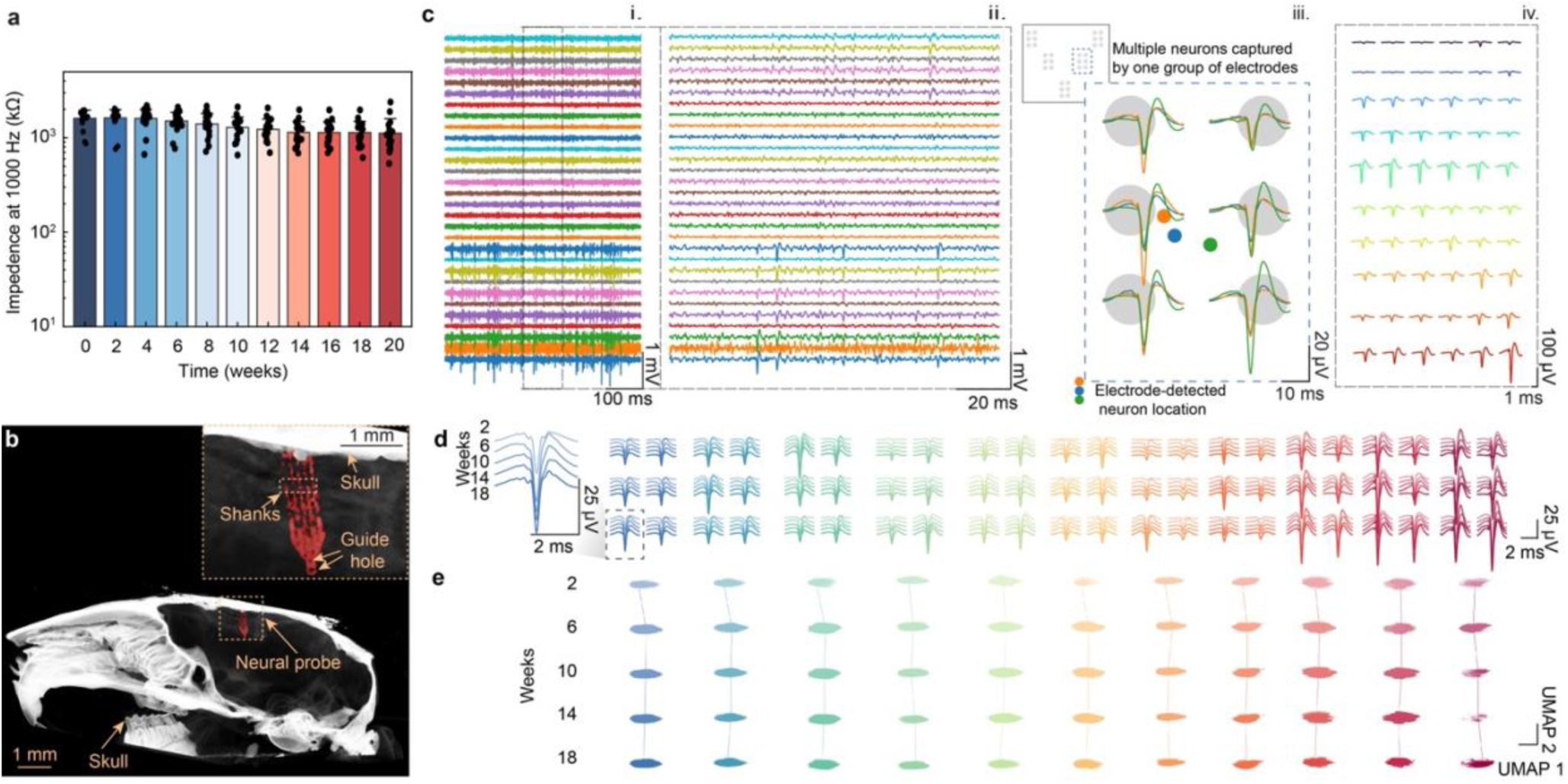
Long-term stable brain electrophysiological recordings across 126 days by FlexiSoft neural probes. a, Dot and bar plots showing time-dependent impedance measurements of FlexiSoft electrodes at 1 kHz in 1× phosphate-buffered saline (PBS) over 20 weeks. Data are mean ± s.d. (n = 16 electrodes). **b,** Micro-computed tomography (Micro-CT) image of an implanted FlexiSoft probe with the inset showing electrodes unfolded inside a mouse brain. **c,** Representative neural recordings from a FlexiSoft probe implanted in a mouse for four weeks. i, Representative raw voltage traces recorded from multiple channels, showing neural activity over a 500 ms window. ii, Zoomed-in view of a 100-ms segment highlighting spike events across different channels. iii, Overlaid spike waveforms detected from a representative electrode array, with spatial positions mapped onto the electrode array (inset). The colored dots indicate the estimated neuron locations. iv, Waveforms of a putative unit simultaneously captured by six electrodes (one electrode array). Each row represents a unit recorded by six electrodes. The displayed waveforms are from representative units recorded across different electrode arrays spanning five shanks of the device, illustrating the distinct amplitude fingerprints that enable single-unit identification. **d,** Average single-unit waveforms recorded at each electrode from a representative FlexiSoft probe over 126 days of recording in a mouse brain. Waveforms from different weeks are color-coded in a gradient. **e,** Time evolution of single-unit waveform features from the same neuron in (**d**), visualized using uniform manifold approximation and projection (UMAP).

We assessed signal stability by three independent analyses in the representative mouse. First, using uniform manifold approximation and projection (UMAP)^33^ (Fig. 3e), we confirmed that spike waveform clusters remained stable over time. Second, we quantified waveform features (Extended Data Fig. 5a-f), including half-width duration, repolarization slope, peak-trough duration, peak-trough ratio, and recovery slope, which remained stable over 126 days (*n* = 22 neurons; *p* > 0.05, one-way ANOVA, Tukey’s multiple-comparisons test; Methods). Third, statistical analysis of waveform similarity, firing rate, and signal-to-noise ratio (SNR) showed no significant changes over the 126 days (Extended Data Fig. 5g-i; *n* = 22 neurons; *p* > 0.05, one-way ANOVA, Tukey’s multiple-comparisons test). To further confirm the stable tracking of the signals from the same neuron over time, we performed additional spatial tracking analyses. The displacement of centroid positions of each neuron defined by their waveform amplitude distribution across multiple nearby electrodes remained < 10 μm over the 126 days period (Extended Data Fig. 5j). Second, the average distance between centroid of different neurons (*r*across) was significantly greater than the displacement of the same neuron (*r*within) over time (Extended Data Fig. 5k-n). The horizontal (4.9 ± 1.9 μm, mean ± s.d.) and vertical (4.3 ± 3.6 μm, mean ± s.d.) displacements of neuronal centroid between weeks 2 (day 14) and 18 (day 126) were smaller than the size of a single neuron soma (Extended Data Fig. 5o). Together, these results confirm that robust FlexiSoft probe enables long-term, stable tracking of same-neuron activity for at least 126 days *in vivo*.

Tracking neural activity from the same neurons across multiple brain regions is crucial for understanding how distributed brain networks communicate and coordinate complex functions such as learning, memory, and motor control. However, current tissue-level flexible probes lack the mechanical robustness required for high-yield implantation while ensuring stable long-term single-unit tracking across distributed brain regions. Here, we investigated whether the enhanced robustness of FlexiSoft probes enables both high-yield implantation and stable tracking of the same neurons across multiple brain regions. To evaluate this, we implanted two 40-channel FlexiSoft probes on one device in the HPC and M1 (Fig. 4a, Methods). Micro-CT imaging of a representative mouse confirmed successful implantation in both regions, approximately 3 mm apart, in a parallel anterior-posterior (A/P) orientation, with intact probe structures (Fig. 4b). Across 13 implanted mice, neural signals were successfully recorded from both brain regions in 11 mice, and the remaining 2 mice had signals from only one region, achieving a 92.3% implantation success rate (24 out 26 neural probes functioning across 13 mice). Over a 12-week recording period, we recorded and analyzed activity from 156 neurons in the HPC and 203 neurons in the M1 region (Fig. 4c). Representative raw traces from an 80-channel probe demonstrated stable multi-channel recordings from both regions (Fig. 4d-e). Consistent neural signals were observed in both HPC and M1 over 12 weeks, with overlaid waveforms confirming stable tracking from the same neurons (Fig. 4f-i). Single-unit activities from the same neurons were detected simultaneously across eight electrodes at each site.

**Fig. 4.**
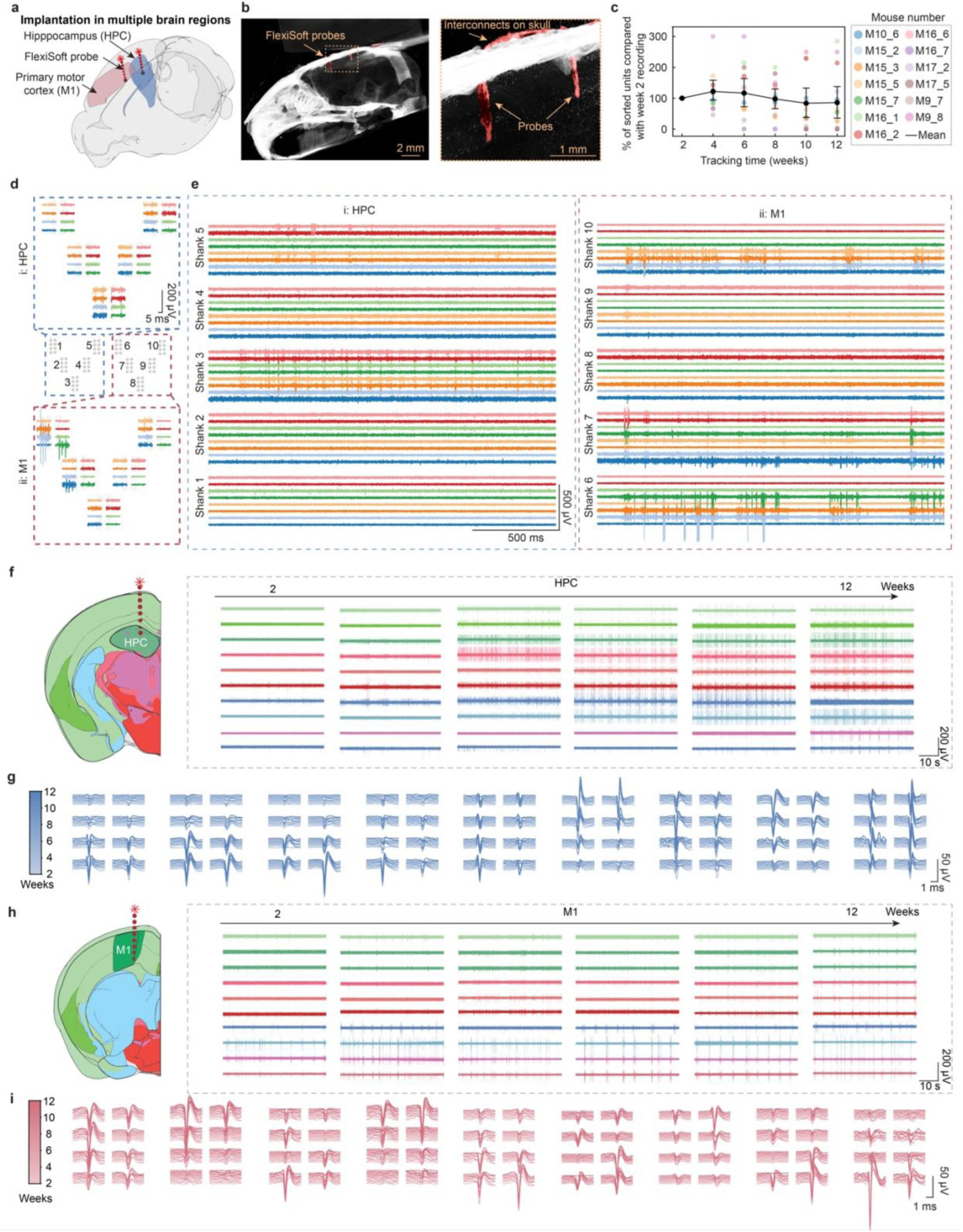
Long-term, stable tracking of activity from the same neurons across multiple brain regions. a, Schematic of multiple region implantation using an 80-channel, multi-shank FlexiSoft probe targeting both the hippocampus (HPC) and primary motor cortex (M1). **b,** Micro-CT image showing two FlexiSoft probes implanted in a single mouse brain. **c,** Tracking yield of units normalized to the initial recording session across 12 weeks. Each point represents a single unit recorded from one mouse at a given session; colors indicate individual mice. In total, 156 units from HPC and 203 units from M1 across 13 mice were analyzed. **d,** Electrode layout of an 80- channel FlexiSoft probe with 40 channels in HPC and 40 in M1. **e,** Representative voltage traces recorded two weeks post-implantation. **f-i,** Representative raw traces (**f** and **h**) and averaged single-unit waveforms (**g** and **i**) from the HPC and M1 over 12 weeks. Gradient color plots in **g** and **i** denote waveform evolution over time.

Our waveform stability evaluation showed that the amplitude of waveforms from individual neurons remained consistent throughout the 12-week recording period in both brain regions. Most waveform features exhibited minimal changes over time (Extended Data Fig. 6a-e; *p* > 0.05, one-way ANOVA, Tukey’s multiple-comparisons test; Method). To further confirm the stability of recording across multiple brain regions, we examined the SNR and firing rate change over time (Extended Data Fig. 6f-g). In the HPC (*n* = 12 mice, 156 neurons) and M1 (*n* = 12 mice, 203 neurons), SNR and firing rates remained stable in 95.8% and 97.3% of neurons, respectively. Together, these findings illustrate that the neural signals remain consistent for 12 weeks of long-term recordings from same neuron at single-cell resolution in both brain regions.

### Long-term stable tracking from same neuron across the entire motor learning task

Learning is a gradual, multi-stage process^34^, with region-specific adaptations emerging at different times. Therefore, tracking the same neuron over extended periods is crucial for understanding how neural activity evolves and interacts throughout these phases. However, existing technologies are limited. They either track neuronal populations in cortical regions for months without addressing the activity from the same neurons^1^ or track a few neurons for only a few days to weeks^35,36^. None of them capture long-term, cross-regional, single-cell dynamics. For example, previous studies have demonstrated dynamic HPC-M1 coupling^37^ at the population level and short-term (one week) activity changes in the same neuron within individual brain regions^1,38^ during motor learning. Yet, how individual neurons adapt over extended learning tasks across both HPC and M1 remains largely unknown. The ability of FlexiSoft probes to provide stable, long-term tracking of the same neurons across multiple brain regions presents a unique opportunity to observe how single-neuron dynamics evolve with learning. Using FlexiSoft, we designed a skilled motor learning task to monitor long-term neural dynamics in both HPC and M1 at the single-neuron and population levels (Fig. 5a).

**Fig. 5.**
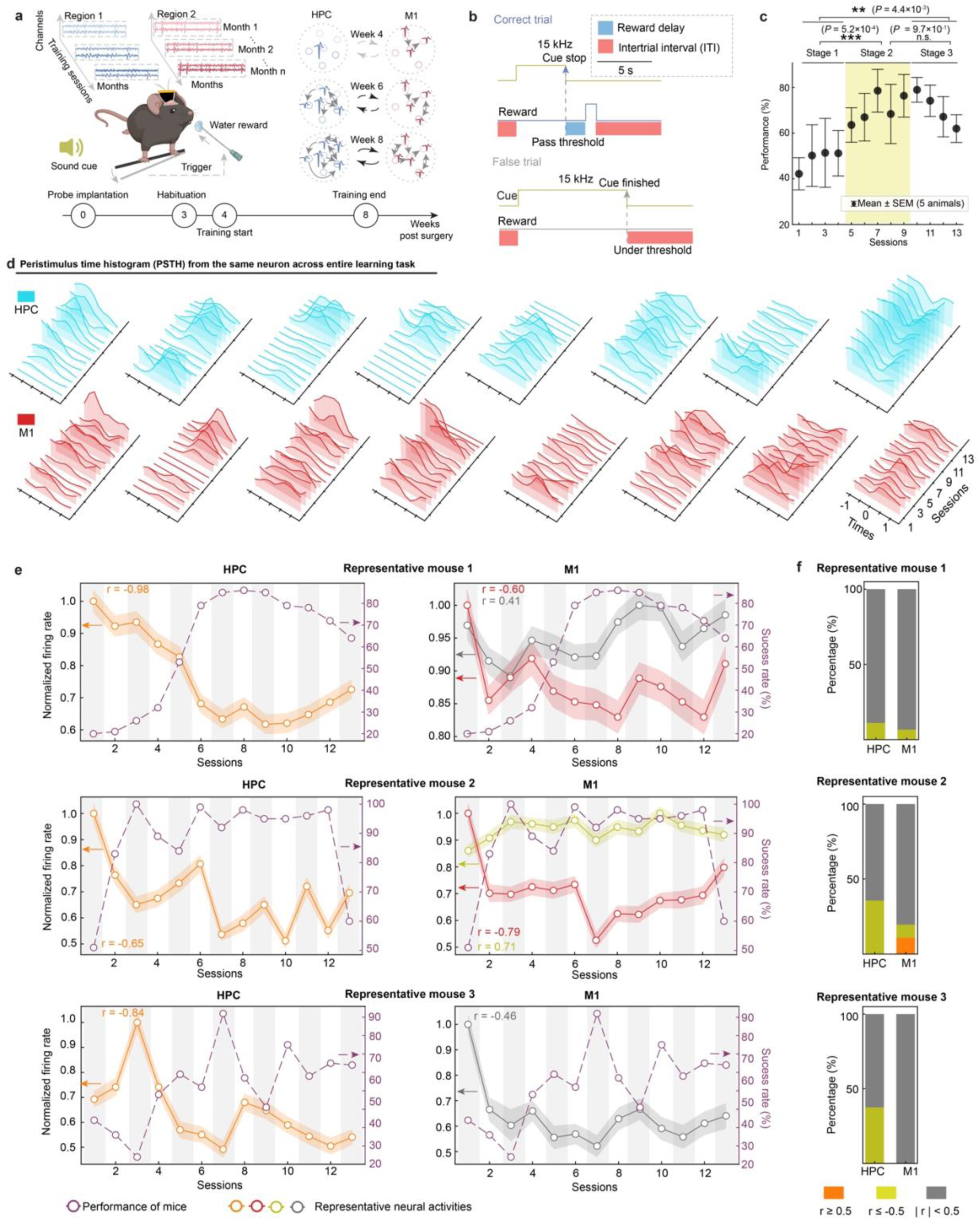
Long-term, stable tracking of single-unit activity from the same neurons across HPC and M1 during a motor learning task. a, Schematic of the behavior paradigm and recording setup. Mice were implanted with FlexiSoft probes in both HPC and M1 and trained to perform a sound cue-triggered joystick task across 13 sessions. **b,** Diagram of the trial structure. A 10-second sound cue initiated each trial. Joystick movements surpass a predefined threshold within the cue window were rewarded with water. **c,** Behavior performance over 13 sessions (n = 5 mice), measured as success rate per session. **d,** Representative single-unit activity from the same neurons in HPC (blue) and M1 (red) across all sessions, aligned to sound cue onset (t = 0). **e,** Session-wide evolution of normalized firing rate from representative neurons (within 1 s after the sound cue) and overlaid with the corresponding mouse performance. Shaded regions indicate mean±SEM of 100 trials per session; Pearson’s r indicates correlation between firing rate and behavioral performance. **f,** Distribution of Pearson’s r between firing rate and behavioral performance across neurons in HPC and M1 for three representative mice. Neurons were categorized into positive correlation (r ≥ 0.5, orange), weak/no correlation (-0.5 < r < 0.5, gray), and negative correlation (r ≤ -0.5, yellow).

We trained mice implanted with FlexiSoft probes in both HPC and M1 to respond to reward-predictive cues over 13 training sessions across one month (Extended Data Fig. 7a-c; Supplementary Discussion 4; Methods). In this task, mice learned to associate joystick movements—either pushing or pulling beyond a displacement threshold—within a sound-cued reaction window to receive a water reward (Fig. 5b). Behavioral analysis revealed significant improvement in the mice performance after training. The success rate in trained mice (*n* = 5 mice) increased significantly between early stages (Session 1-4) and later stages (Sessions 5-13) (Fig. 5c, Supplementary Video 1; two-tailed, unpaired t-test), consistent with patterns observed in previous motor learning paradigms^39,40^. Based on performance metrics, we categorized the learning process into three stages^41,42^: Stage 1 represents the early learning phase, characterized by a low success rate (49 ± 5%, mean ± SEM) and a long reaction time to hit the joystick following the sound cue (2.99 ± 0.09 s, mean ± SEM; Extended Data Fig. 7d). Stage 2 corresponds to the consolidation phase where performance improves (71 ± 4%, mean ± SEM) and reaction time decreases significantly (2.55 ± 0.07 s, mean ± SEM) compared to Stage 1. Stage 3 represents the overtraining phase, where performance slightly declines, with the average success rate plateauing (71 ± 4%, mean ± SEM) and reaction time increases marginally, though not significantly (2.65 ± 0.08 s, mean ± SEM). These results indicate that the mice developed a learned association between joystick movement and reward in response to the sound cue.

We then investigated how neural dynamics evolved throughout the training period. We tracked 69 neurons in the HPC and 54 neurons in the M1 (*n* = 5 mice) and analyzed their firing activity 1 s before and after the sound cue during successful trials across all sessions. We identified the neuronal subtypes across HPC and M1 based on their dynamic response to the learning. For example, we observed neuronal subtypes that either exhibited significant firing rate increase/decrease across learning sessions (especially Stage 1) or remained stable throughout the learning (Fig. 5d; Extended Data Fig. 7e-f). To further link intrinsic single-cell dynamics with behavior performance, we calculated Pearson correlation coefficients between firing rate and task success rates, classifying neurons as negatively correlated (*r* ≤ -0.5), weak or no correlation (-0.5 < *r* < 0.5), or positively correlated (*r* ≥ 0.5). Notably, we found neurons strongly negatively correlated with task performance. For example, one representative HPC neuron exhibited *r* = -0.98, with firing rate decreasing as task success rate improved (Fig. 5e). Across mice, negatively correlated neurons (*r* ≤ -0.5) were observed in both HPC and M1, with a greater proportion in HPC. In contrast, positively correlated neurons (*r* ≥ 0.5) were only found in M1 and represented a smaller fraction of neurons. Most neurons in both regions exhibited weak or no correlation (-0.5 < *r* < 0.5; Fig. 5f).

In addition, heatmaps of normalized neural activity aligned to sound cue onset revealed similar temporal dynamics and cue responses across HPC and M1 neurons (Extended Data Fig. 8a). Notably, many neurons exhibited dynamic changes during early sessions and stabilized during later stages, consistent with previously reported patterns of neural activity at the population level during motor learning^37,43,44^ and short-term single-neuron tracking results over a few days^2^.

To further investigate how overall neural dynamics evolved in the HPC and M1, we applied Gaussian-process factor analysis^45^ (GPFA), a trial-based linear dimensionality reduction method to extract low-dimensional neural trajectories. This approach allowed us to examine time-varying latent neural dynamics across sessions within a shared low-dimensional manifold. The results show that GPFA neural trajectories from the HPC exhibited greater variability during Stage 1 and Stage 2 compared to Stage 3, as evidenced by changes in GPFA trajectory shape and orientation (Extended Data Fig. 8b). In comparison, the neural trajectory in M1 remained relatively stable, with a nearly unchanged shape and orientation. In addition, the top two latent dimensions showed significant changes during Stage 1 in both regions but converged to more consistent, stable dynamics by Stage 3 (Extended Data Fig. 8c-d).

In summary, these findings demonstrate that FlexiSoft enables stable, long-term tracking of activity of the same neurons across extended learning periods and multiple brain regions. This capability allowed us to classify distinct neuronal subpopulations based on their relationship to task performance—identifying specific neurons with positive, negative, or weak/no correlation to task performance. In contrast to the conventional methods limited to population-level changes, FlexiSoft probes provide insights into how individual neurons contribute to different phases of learning. While most changes in firing rates occurred during the early learning phase (Sessions 1- 4), both single-neuron activity and network-level dynamics stabilized by the overtraining phase (Sessions 10-13), reflecting a transition from early neural adaptation to sustained task performance.

### FlexiSoft for customizable curved-path implantation and deep implantation

The unique combination of robustness and tissue-level flexibility of FlexiSoft probes enables a new class of neural interface strategies. One key advantage is their ability to support curved-path implantation, where the probe is inserted along a curved or serpentine trajectory to reach targeted brain regions. This approach is especially critical when the target site lies beneath sensitive brain areas; by following a curved trajectory, the probe can bypass and preserve overlying tissues, avoiding the damage that conventional straight-line insertion would risk. Traditional rigid probes lack the flexibility to follow such curved trajectory, while conventional flexible probes often fail to maintain mechanical integrity during curved implantation.

We demonstrated that the FlexiSoft probe can be readily implanted along a curved or serpentine trajectory through a curved shuttle, enabling several important advantages: (i) avoiding damage to sensitive brain regions (Fig. 6a); (ii) minimizing disruption to critical blood vessels (Fig. 6b); and (iii) enable signal recording along curved/serpentine trajectories that would be inaccessible through a single vertical insertion. *In vitro* testing using brain-mimicking agarose gel (Extended Data Fig. 9a-b) confirmed that FlexiSoft probe could be implanted stepwise along a curved trajectory while maintaining structural integrity. Micro-CT imaging further demonstrated successful curved-path implantation *in vivo* within a mouse brain (Fig. 6c-d). Unlike traditional implants, which require a uni-directional trajectory perpendicular to the skull, FlexiSoft probes can follow an almost parallel trajectory relative to the skull surface (Fig. 6e, Extended Data Fig. 9c-d). Importantly, neural recordings along the curved trajectory remained stable over time, as shown by representative raster plots and waveform analyses 2 weeks post-implantation (Fig. 6f- g).

**Fig. 6.**
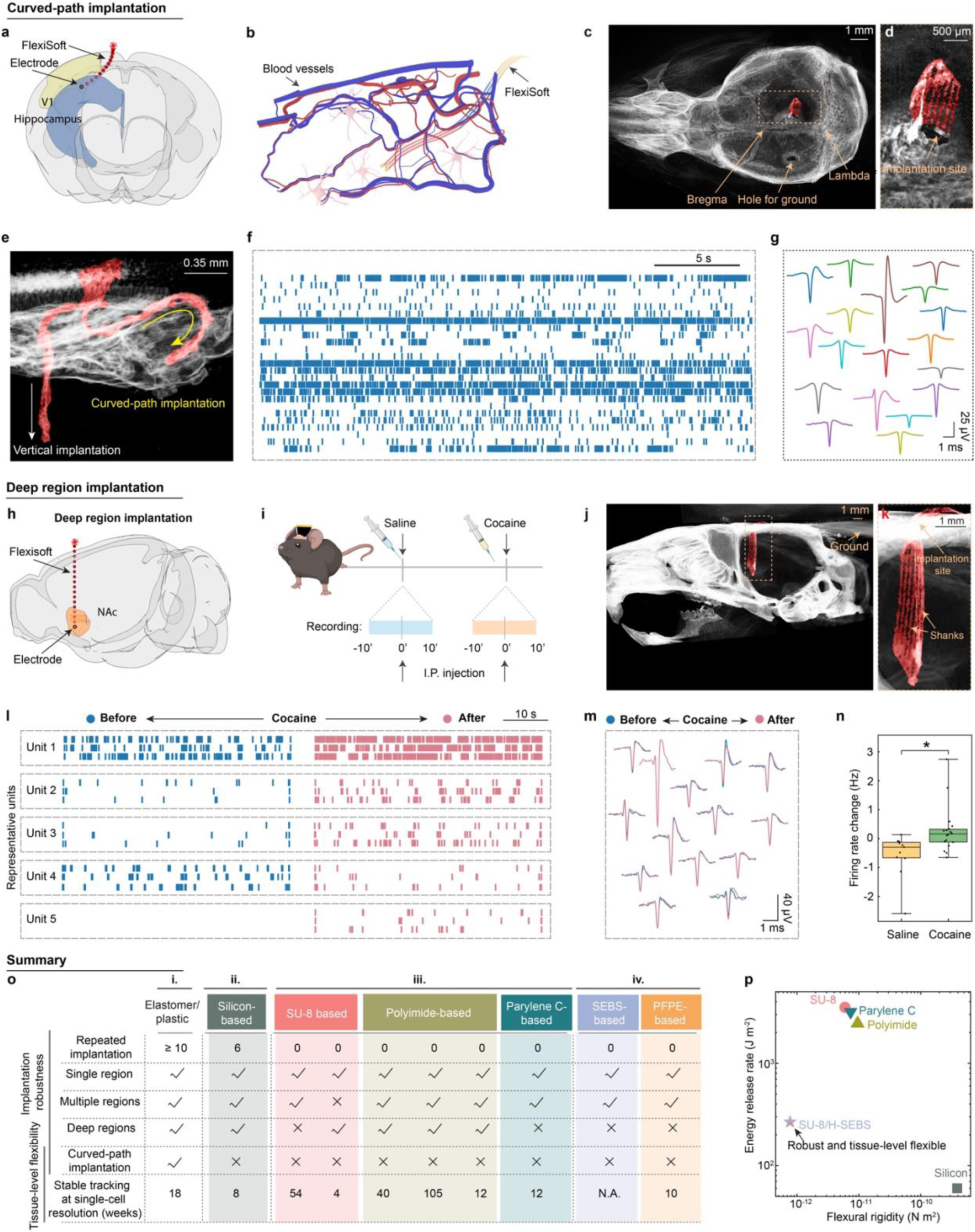
FlexiSoft probe-enabled curved-path and deep brain region implantations. a, Schematic of curved-path implantation bypassing the overlying brain area to reach deep target region. **b,** Schematic illustrating curved-path implantation designed to avoid major blood vessels. **c,** Micro-CT image showing curved-path implantation in a mouse brain from a ventral-to-dorsal view. **d,** Zoomed-in image of (**c**) showing the FlexiSoft probe following a curved path implantation. **e,** Micro-CT image from a lateral-to-medial view showing simultaneous vertical and curved-path implantation in the same mouse brain. **f-g,** Representative spike raster plot (**f**) and single-unit waveforms (**g**) recorded two weeks after curved-path implantation. **h,** Schematic of deep brain implantation targeting the nucleus accumbens (NAc). **i,** Schematic of cocaine administration experiment. **j,** Micro-CT image showing a FlexiSoft probe implanted in a deep mouse brain region. **k,** Zoomed-in view of the implantation site from the orange dashed box in (**j**). **l-m,** Representative raster plots (**l**) and waveforms (**m**) from the same neurons before and after cocaine administration. **n,** Box plot of the firing rate change after saline and cocaine administration. **o,** Comparative matrix of existing neural probes^7,16,17,47–51^, showing FlexiSoft probes uniquely combine robustness and flexibility across all implantation scenarios, including repeated implantation-retrieval, multi- region targeting, and curved-path trajectories. **p,** Energy release rate versus flexural rigidity for rigid, flexible and FlexiSoft probes. All samples are under identical tensile load (constant far-field uniaxial stress, *σ* = 50 MPa).

Another key advantage of the FlexiSoft probe is its ability to target deep brain regions. As a demonstration, we implanted FlexiSoft probes into the mouse nucleus accumbens (NAc) to monitor neural responses to drug interventions (Fig. 6h-i). Micro-CT imaging confirmed successful implantation to depths of 4-5 mm without any structural damage (Fig. 6j-k). We recorded neural activity before and after the administration of saline and cocaine, enabling tracking of drug-induced changes in neural dynamics at single-neuron resolution. Representative raster plots showed increased activity following cocaine administration, while waveform shapes remained stable over time (Fig. 6l-m). Statistical analysis further confirmed significant differences in firing rates between saline and cocaine groups (Fig. 6n).

Together, these results demonstrate FlexiSoft’s robust and flexible performance across a wide range of implantation paradigms, including both curved-path and deep brain region implantations.

## Discussion

We introduced the FlexiSoft platform, a heterostructure neural probe that integrates flexible plastic with a soft elastomer layer (Fig. 6o-p). Notably, the FlexiSoft platform achieves exceptional mechanical robustness while maintaining tissue-level flexibility, reducing energy release rates by 13.8 times and flexural rigidity by 7.7 times compared to traditional flexible electronics of the same thickness. This enhanced robustness enables repeated implantation-retrieval cycles, curved- path implantation, and deep brain region implantation. Meanwhile, the preserved tissue-level flexibility supports chronic stable implantation, as evidenced by a minimized brain immune response to FlexiSoft probes (Extended Data Fig. 10). Fluorescence intensity measurements of neurons, astrocytes, and microglia around the implant were comparable to those observed with ultrathin SU-8 flexible probes, indicating good long-term biocompatibility. Together, these features allow for stable tracking of activity from the same neurons for up to 126 days (Fig. 3).

The combined mechanical robustness and tissue-level flexibility of FlexiSoft probes enables stable tracking of activity from the same neurons simultaneously across multiple brain regions. Using this approach, we tracked 156 neurons in HPC and 203 neurons in M1 region over 12 weeks across 13 mice (Fig. 4), and during a motor learning task, which represents, to our knowledge, the first demonstration of dynamic single-neuron tracking across these regions during motor learning (Fig. 5). This ability allowed us to identify distinct neuronal subtypes based on their long-term firing dynamics during motor learning: some neurons exhibited flexible, early-phase adaptations, while others maintained stable activity across all learning stages. Furthermore, stable single-cell tracking uncovered functional specialization within and across brain regions: in HPC, neurons with firing rates negatively correlated with task performance likely support early flexible encoding that diminishes with learning, whereas in M1, neurons with positive correlations appear to contribute to the consolidation and execution of learned motor patterns. Together, these findings bridge the gap between population-level studies and single-neuron tracking, revealing how functional specialization across brain regions during skill acquisition.

The FlexiSoft platform supports versatile material combinations beyond SU-8 and H-SEBS, such as SU-8/PFPE-DMA, achieving lower energy release rates compared to homogeneous flexible structures such as polyimide (Fig. 6p and Supplementary Fig. 2). This material tunability enhances the design space for FlexiSoft BCIs, constrained only by available fabrication methods.

Looking ahead, centimeter-scale, mechanically robust FlexiSoft probes with repeated implantation-removal capability can be readily translated into clinical applications, enabling stable, long-term brain-computer interfaces in humans. Their robustness and flexibility make them ideal for deep, multi-region, and curved-path brain implantations in chronic neurophysiological and medical applications.

## Methods

### 1. Materials and electronics

#### Materials

H-SEBS (Tuftec H1062) was obtained from Asahi Kasei. SU-8 2002 photoresist and all other photoresists and developers in the nanofabrication were obtained from MicroChem Corporation unless otherwise specified.

#### FlexiSoft fabrication

FlexiSoft fabrication begins with cleaning a 3-inch thermal oxide silicon wafer (University Wafer). The wafer is sequentially rinsed with acetone, isopropanol (IPA), and water, followed by drying with a nitrogen stream and exposure to 100 W, oxygen (O2) plasma (40 sccm O2) for 5 min. A 100-nm-thick nickel (Ni) sacrificial layer is then patterned on the cleaned silicon wafer. The photoresist (PR) pattern for the nickel layer is prepared as follows: HMDS is spin-coated at 2000 rpm for 30 s, followed by LOR 3A at 2000 rpm for 30 s following a hard-bake at 180°C for 5 min. PR S1805 is then spin-coated at 2000 rpm for 30 s and hard-baked at 115°C for 1 min. The photoresists are exposed to 40 mJ/cm² UV light and developed in CD-26 for 1 min, rinsed with deionized (DI) water for 30 s, and blown dry. Following PR patterning, a 100-nm Ni layer is thermally deposited on the wafer (Sharon) at a rate of 1 Å/s and lifted off in Remover PG for 3 hours. Next, the bottom SU-8 encapsulation layer is patterned. SU-8 2002 is spin-coated at 3000 rpm for 30 s and pre-baked at 60°C for 1 min, then at 95°C for 1 min. SU-8 is exposed to 100 mJ/cm² UV light and post-baked at 60°C for 1 min, followed by 95°C for 1 min. The SU-8 is developed in SU-8 developer for 90 s, rinsed with IPA for 30 s, and blown dry. A final hard bake at 180°C for 1 hour completes this step. The third step is patterning the gold (Au) cable layer using lift-off process (same for other metal layer patterning). A 5/60/5 nm Cr/Au/Cr layer is deposited onto the bottom SU-8 encapsulation layer using an electron-beam evaporator (Denton) with a PR pattern and lifted off in Remover PG for 5 hours. The fourth step involves depositing the platinum (Pt) layer. A 3/40 nm Cr/Pt layer is patterned and deposited by an electron-beam evaporator (Denton) and lifted off in Remover PG for 2 hours. The fifth step is preparing the top SU-8 encapsulation layer over the Au cable layer. The fabrication process for the top SU-8 layer mirrors that of the bottom SU-8 layer. The final step is patterning the H-SEBS top supporting layer. H- SEBS is spin-coated at 2,000 rpm for 45 s and baked at 80°C for 1 hour. A 3/40 nm Cr/Au layer is deposited on the H-SEBS. S1822 is then spin-coated on top of the H-SEBS at 2,000 rpm for 45 s, followed by a 3-min bake at 80°C. The photoresist is exposed at 310 mJ/cm² to form the pattern and developed in MF 319 for 1 min, rinsed with DI water for 30 s, and blown dry. The wafer is subsequently etched in Au and Cr etchants for 40 s and 20 s, respectively, to form the Au hard mask pattern for the final oxygen plasma etching (ICP power: 1500 W, forward plasma power: 300 W, O2 flow: 40 sccm, duration: 15 min; Oxford PlasmaPro 100 Cobra 300).

#### Flip-chip bonding and releasing of FlexiSoft

The flexible flat cable (Molex or customized in PCB Way) is bonded to the input/output pads of the electronics using a flip-chip bonder (Finetech Fineplacer). To encapsulate the I/O bonding region, polydimethylsiloxane (PDMS, DOW 1696157 SYLGARD 170 Silicon Elastomer) is applied. For probe release, the electronics are immersed in a nickel (Ni) etchant solution (TFB, Transene Company) at room temperature for 3-4 hours. After complete release from the substrate, the Ni etchant is replaced with deionized (DI) water to thoroughly cleanse the probe. Before surgery, the electronics are first placed in IPA for 30 min for disinfection and then in Poly-D-Lysine for modification to enhance the adhesion of neurons. Prior to implantation, the anchors of the electrodes were cut, and the electrodes were fully detached from the surface of the silicon wafer.

### 2. Characterization

#### Thickness measurements

All thickness measurements were carried out using a DektakXT stylus profiler (Bruker). The force applied was set to 3 mg and the scan speed was 0.67 μm s^−1^. Two-point surface levelling was applied using the commercial software of the tool.

#### SEM imaging and EDS element analysis

A metal sputter coater was used to deposit a 20-nm-thick Pt/Pd layer and reduce the charging under the electron beam during the SEM imaging process. SEM images were taken under 5 kV voltage using a JEOL 7900F SEM instrument. Before FIB milling, another 100-nm-thick layer of Pt was deposited using the metal sputter coater atop the sample to prevent damage of the area of interest under ion beams. The FIB process followed conventional procedures using an FEI Helios 660 instrument in which a 30 kV Ga beam was used for rough milling and a reduced 10 kV voltage was used for fine milling. To inspect the sample surface of the FIB-milled cross-section, a secondary electron detector under 3 kV voltage was used to take the SEM images, which showed clear contrast and distinct layering out of the samples.

#### *In vitro* impedance measurement and analysis of FlexiSoft

Before surgeries and during repeated implantation test, the impedance of the electronics was tested using an Intan 1024ch recording controller. The electronics were immersed in a 1× PBS solution (VWR International, 97063-660) with the flexible cable connected to the Intan controller. A stainless-steel electrode submerged in the PBS solution serves as the reference. For repeated implantation impedance measurement, yield was defined as the percentage of functional channels, where functionality was determined by impedance values falling between 50 kΩ and 5000 kΩ at 1 kHz. Channels that failed this criterion at any timepoint were considered permanently non-functional and excluded from subsequent analyses.

### 3. Animal experiments

#### Animals and ethical compliance

Mature adult male C57BL/6 mice, 16 weeks of age (Charles River Laboratories), were used throughout this study. The mice were maintained at 22 ± 1 °C with humidity ranging from 30% to 70% and on a 12-hour light/dark cycle and were fed ad libitum. All procedures complied with the National Institutes of Health guidelines for the care and use of laboratory animals and were approved by the Harvard University Institutional Animal Care and Use Committee under protocols # 20-05-368-1.

#### Stereotaxis implantation

During intracranial implantation surgery, mice were anesthetized using 2–3% isoflurane and maintained under 0.75–1% isoflurane. Stainless steel screws were inserted into the cerebellum to serve as ground electrodes. A 2 × 2 mm craniotomy was performed to expose the cortical surface, followed by removal of the dura mater. The FlexiSoft neural probes, attached to a customized tungsten shuttle (50 μm in diameter), were secured to a micromanipulator on a stereotaxic frame. This micromanipulator was used to carefully insert the electronics into the brain to the desired depth. The craniotomy was then sealed with a silicone elastomer (World Precision Instruments), and ceramic bone anchor screws and dental methacrylate were used to secure the flat flex connector (FFC) and electrode set onto the mouse’s skull. For centimeter-long FlexiSoft implantation, a customized film-like shuttle (10 cm long, 500 µm wide, 30 µm thick) was used. For curved-path implantation, a customized bent tungsten shuttle (50 μm in diameter) was employed.

#### Electrophysiological data acquisition

Implanted mice were head-fixed using a custom 3D-printed setup. The flexible flat cables were connected to a customized PCB, which was then interfaced with either a Blackrock or Intan headstage. Neural signals were subsequently recorded using the Blackrock recording system (CerePlex Direct) and Intan RHD recording system (Intan 1024ch Recording Controller). The setup was placed on an optic table and covered by a Faraday cage.

#### Joystick training

Water restriction began three days prior to the training tasks. The mice were trained in the joystick- moving task for 13 sessions across 1 month. Intan recordings on the implanted mice were performed during the training sessions. The setup was placed on an optic table and covered by a Faraday cage. The behavior setup was controlled by a microcontroller. Each training session consists of 100 repeated trials. Each trial begins with a 10-second 15-kHz tone, during which a joystick movement passing a threshold of 1.5 mm would be rewarded (delayed for 1 s after trigger) with water (∼8 µl per trial) and stops the tone playing, followed by an intertrial interval of 6s.

#### Micro-CT

Mice implanted with single FlexiSoft probe, multiple FlexiSoft probes, curved-path FlexiSoft and deep FlexiSoft were sacrificed and decapitated in accordance with the guidelines approved by Harvard University’s Animal Care and Use Committee. Following decapitation, the mouse head was imaged using an HMXST Micro-CT X-ray scanning system (model HMXST225; Nikon Metrology, Inc.). The scanning parameters were set to 100 kV and 87 μA. Prior to scanning, shading correction and flux normalization were performed to optimize the X-ray detector. Micro-CT sinograms were calibrated for centers of rotation and reconstructed into 2D images using CT Pro-3D software (ver. 2.2; Nikon-Metris). The reconstructed images were then processed for 3D rendering and analysis using VGStudio MAX software (ver. 3.1; Volume Graphics GmbH).

#### Immunostaining and corresponding image data analysis

Immunohistochemistry and confocal fluorescence imaging were carried out following standard protocols. Mice were euthanized with sodium pentobarbital (40–50 mg/kg) and perfused transcardially with 30-40 ml of 1× PBS, followed by 40 ml of 4% paraformaldehyde (PFA). The brains, implanted with SU-8/H-SEBS probes, were extracted, post-fixed in 4% PFA at 4 °C for 24 h, and then sequentially immersed in sucrose solutions with gradient concentrations (10%, 20%, 30%, w/v) until they sank. The samples were embedded in optimal cutting temperature compound and sectioned at 30 μm using a cryostat. 0.3% Triton X100 in 1× PBS was used for permeabilization for 30 min, following a 2 hr-blocking step using 2% Bovine Serum Albumin (BSA) in PBST. The sections were incubated overnight at 4 °C with primary antibodies (Anti-NeuN: 1:500, Abcam ab104224; Anti-GFAP: 1:500, Abcam ab4674; Anti-IBA1: 1:500, Abcam ab178846). After three times PBST (0.1% Triton X100 in 1× PBS) washes, secondary antibodies were applied at 4 °C overnight (Goat anti-Mouse IgG (H+L), Alexa Fluor™ 647: 1:1000, ThermoFisher A-21235; Goat anti-Chicken IgY (H+L), Alexa Fluor™ 546:1:1000, ThermoFisher A-11040; Goat anti-Rabbit IgG (H+L), Alexa Fluor™ 488:1:1000, ThermoFisher A-11008). DAPI staining (10 μg mL^-1^) was performed for 30 min before 3 times final PBST wash, following brain slice encapsulation by a cover slide using Cytoseal. Imaging was conducted using a ZEISS LSM 900 confocal laser scanning microscope, and images were captured with the ZEN Microscope Software.

Image analysis of all horizontal slices was performed using ImageJ software in combination with a custom Python script. For each image, the shortest distance from every pixel to the boundary of the FlexiSoft electronics was calculated. Pixel intensities were then grouped into bins at 35 μm intervals based on these distances. Within each bin, the mean fluorescence intensity was calculated and subsequently normalized to a baseline, defined as the average intensity of pixels located 500 – 600 μm away from the FlexiSoft boundary. Normalized intensity profiles were averaged across samples, and the standard deviation was calculated to quantify inter-sample variability.

### 4. Mechanical simulation and analysis

#### Finite-element analysis

COMSOL Multiphysics 6.1 was used to analyze the mechanical properties of various neural probes. The objectives of the simulations were as follows: (1) To assess the stress concentration of SU-8 electronics and SU-8/H-SEBS hybrids prior to crack propagation; (2) To evaluate the energy release rate of SU-8 electronics and SU-8/H-SEBS hybrids with varying crack lengths; (3) To examine the changes in energy release rate with variations in thickness and modulus of the soft supporting substrate layer. The 4-μm-thick SU-8 probes consist of a single SU-8 layer. Additionally, for single-layer electronics, we substituted SU-8 with polyimide and silicon to observe changes in energy release rate with crack length. The 8-μm-thick SU-8/H-SEBS probes comprise one 4-μm-thick SU-8 layer and one 4-μm-thick H-SEBS layer. We adjusted H-SEBS thickness and Young’s modulus to observe corresponding changes in the energy release rate. In the energy release rate simulations, we employed five sets of predefined meshes (ultrafine, finer, fine, normal, and coarse) to ensure convergence of results. The mesh layers in the thickness direction are refined and consisted of at least four layers.

### 5. Data analysis

#### Spike sorting and assessment of single-unit stability over time

Electrical recordings processing and sorting were performed through a Python pipeline using MountainSort^32^ and SpikeInterface^46^. Raw recordings were first bandpass filtered for range 300- 3,000Hz. Then common median reference was removed to reduce common-mode noises during recording. To extract spike waveforms, a spike detection threshold, ranging from 3 to 5 times the standard deviation away from the mean dependent on the recording base noise level, was applied. Noisy putative unit candidates were further manually curated. Sorting results visualizations were carried out using Scanpy and UMAP^33^ for the waveform features. For multiple group comparisons, one-way ANOVA followed by Tukey’s post hoc test was used. P-values < 0.05 were considered statistically significant.

### 6. Statistics and reproducibility

#### Data availability

All data and materials that supporting the findings of this study are available within the paper and its Supplementary Information. Source data are provided with this paper.

#### Code availability

The authors declare that all the code supporting the findings of this study are available within the paper and its Supplementary Information.

## Acknowledgments

We thank Liu Group, Dr. Y. Zhou and Dr. H. Zhu for insightful discussions. J. Liu acknowledges the support from the Aramont Fund for Emerging Science Research, Harvard Grid award, and NIH/NICHD 1R01HD115272.

## Author contributions

J.L., N.L., and X.L. conceived the idea. X.L. developed the nanofabrication protocol, fabricated and characterized electronics. X.L. and Z.W. conducted mechanical simulations. X.L. and J.L. performed implantation. X.L. and A.R. recorded signals. J.C. developed animal training set up. J.C., X.Z., A.R. and X.L. performed the animal training. X.Z. developed the spike sorting and curation pipeline. X.Z. and H.Z. performed sorting and curation. X.L. and X.Z. analyzed the data. X.L. scanned micro-CT. H.Y. and X.L. carried out mechanical test. J.J.V. and N.L. provided critical guidance on mechanical simulation and mechanical test. A.L., X.Y. Z. and X. L. performed immunostaining experiments. Y. H., H.S., W.W. and A.A. contributed critical discussions and inputs on the figures. X.L. drafted the manuscript. All authors discussed, commented and provided revision on the manuscript. J.L. supervised the study.

## Additional information

Correspondence and requests for materials should be addressed to Jia Liu.

---

**Extended Data Fig. 1.**
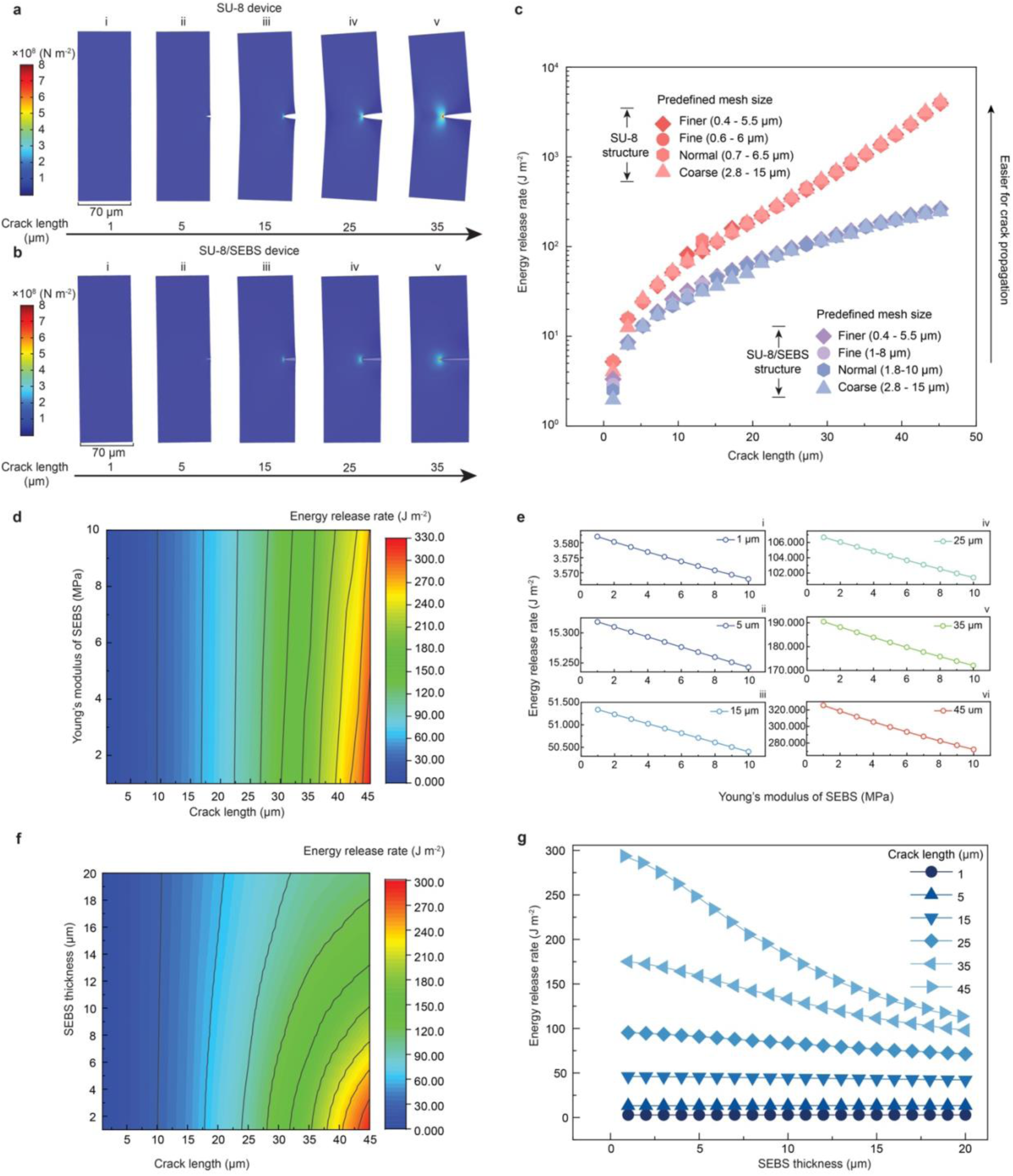
Finite element analysis (FEA) of SU-8 structure and FlexiSoft SU- 8/H-SEBS heterostructure before and after crack propagation. a-b, Stress distribution in an SU-8 structure (4 μm thickness, 70 μm width) (a) and an FlexiSoft heterostructure (4 μm thickness per layer, 70 μm width) (b) for various crack lengths (1, 5, 15, 25, 35 μm) after crack initiation. c, Simulation results showing the energy release rate of the SU-8 and FlexiSoft structures from (a), varying with crack length from 1 to 45 μm. d-g, Impact of H-SEBS substrate thickness, Young’s modulus, and SU-8 flexible layer crack length on energy release rate. Contour map in (d) illustrating the effect of H-SEBS substrate thickness on the energy release rate. Line-cut (e) from (d) showing the energy release rate as a function of H-SEBS Young’s modulus. Contour map in (f) demonstrating the effect of H-SEBS thickness on the energy release rate. Line-cut (g) from (f) showing the energy release rate as a function of H-SEBS thickness.

**Extended Data Fig. 2.**
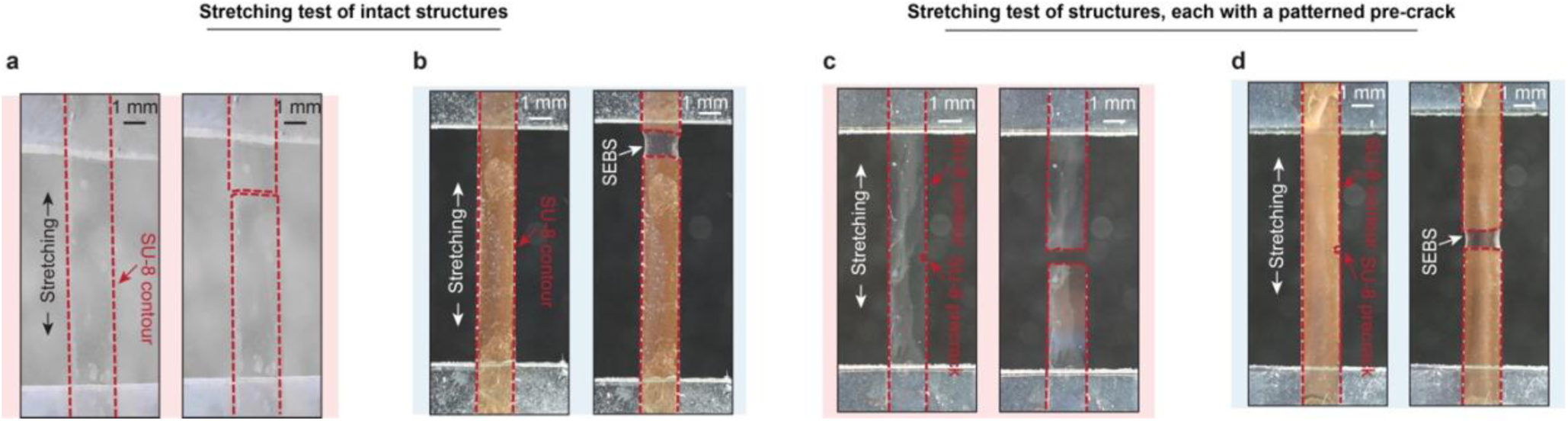
Tensile test for SU-8 and FlexiSoft structures. a-d, Photographs of tensile test without (a-b) or with (c-d) a pre-crack.

**Extended Data Fig. 3.**
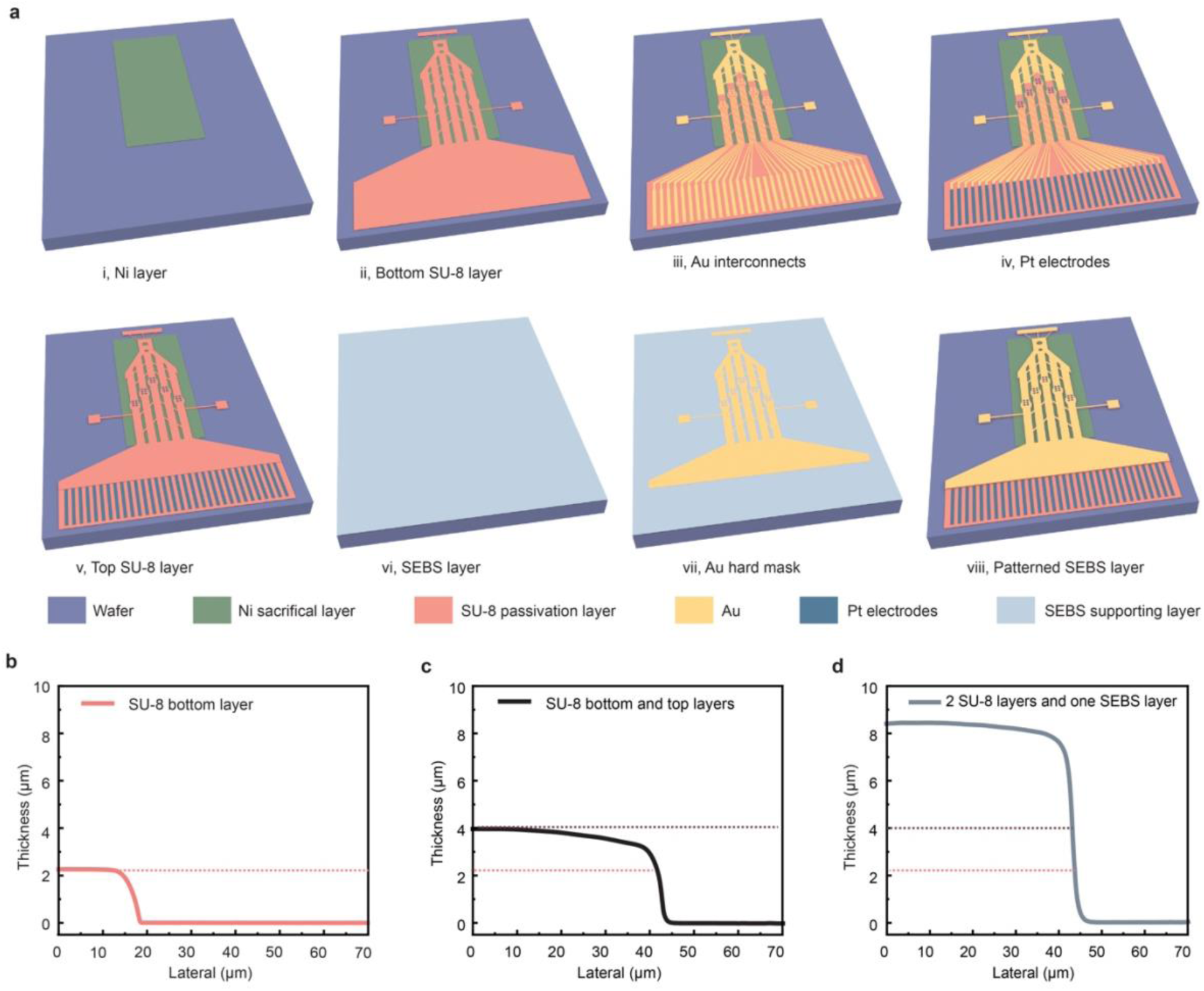
Fabrication of FlexiSoft neural probes. a, Step-by-step schematics of the FlexiSoft probe fabrication process. b-d, Profilometry measurement confirming the thickness of the bottom SU-8 encapsulation layer (b), bottom and top SU-8 encapsulation layers (c), and FlexiSoft structure including both SU-8 layers and H-SEBS soft elastomer layer (d).

**Extended Data Fig. 4.**
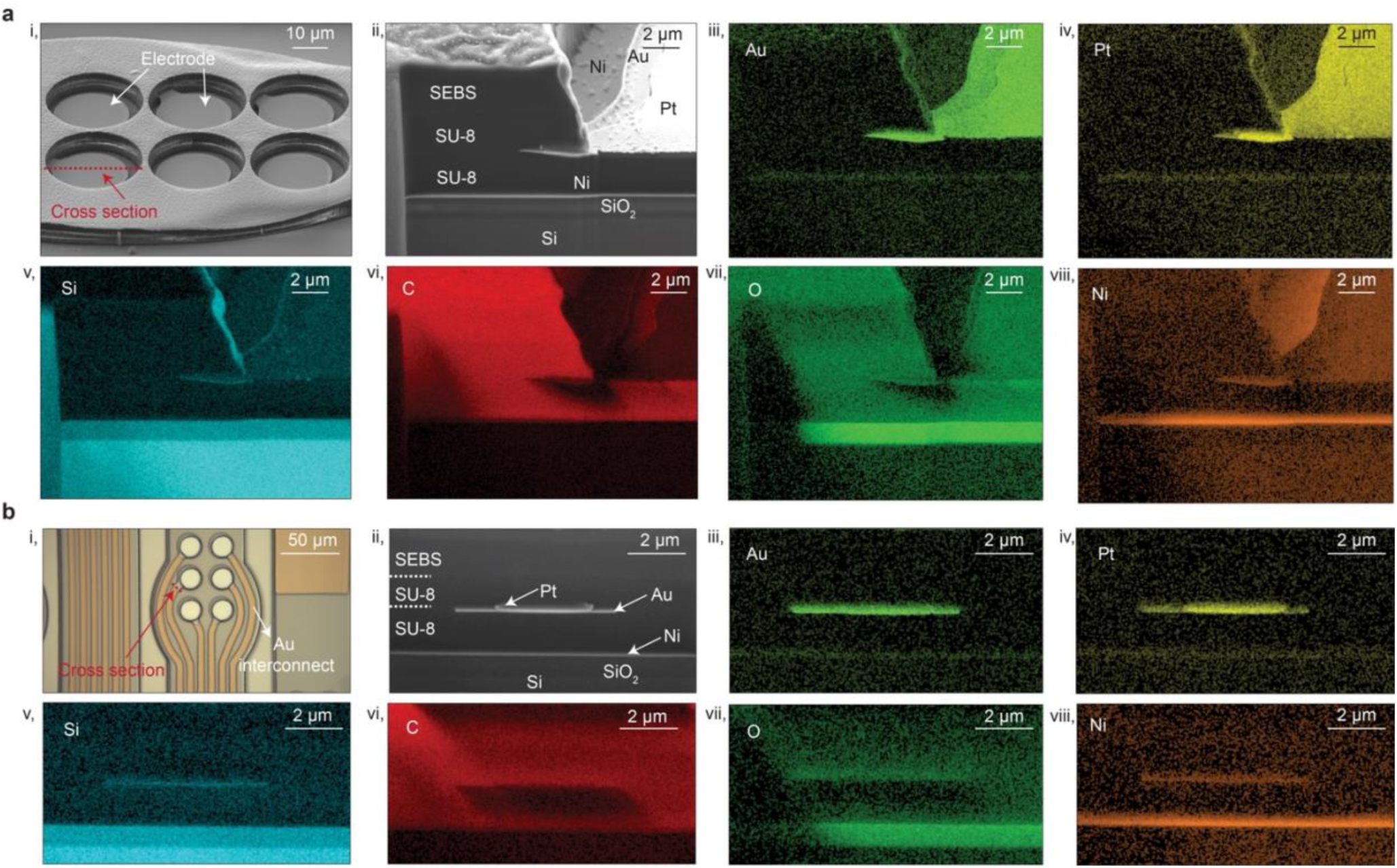
Characterization of FlexiSoft neural probes. a, Cross-sectional structural and element analysis of a representative FlexiSoft structure: i, SEM image of one shank with six electrodes. ii, Focused ion beam (FIB) cross-section of one representative electrode along the red dashed line in (i). iii-viii, Elemental mapping of Au, Pt, Si, Ni, C, and O showing the multilayer structure of the cross-section of (ii). b, Cross-sectional structural and element analysis of a FlexiSoft structure with Au interconnects: i, BF image showing the FIB milling location. ii, Cross-sectional SEM image along the red dashed line in (i). iii-viii, Corresponding elemental mapping of Au, Pt, Si, Ni, C, and O showing the multilayer structure.

**Extended Data Fig. 5.**
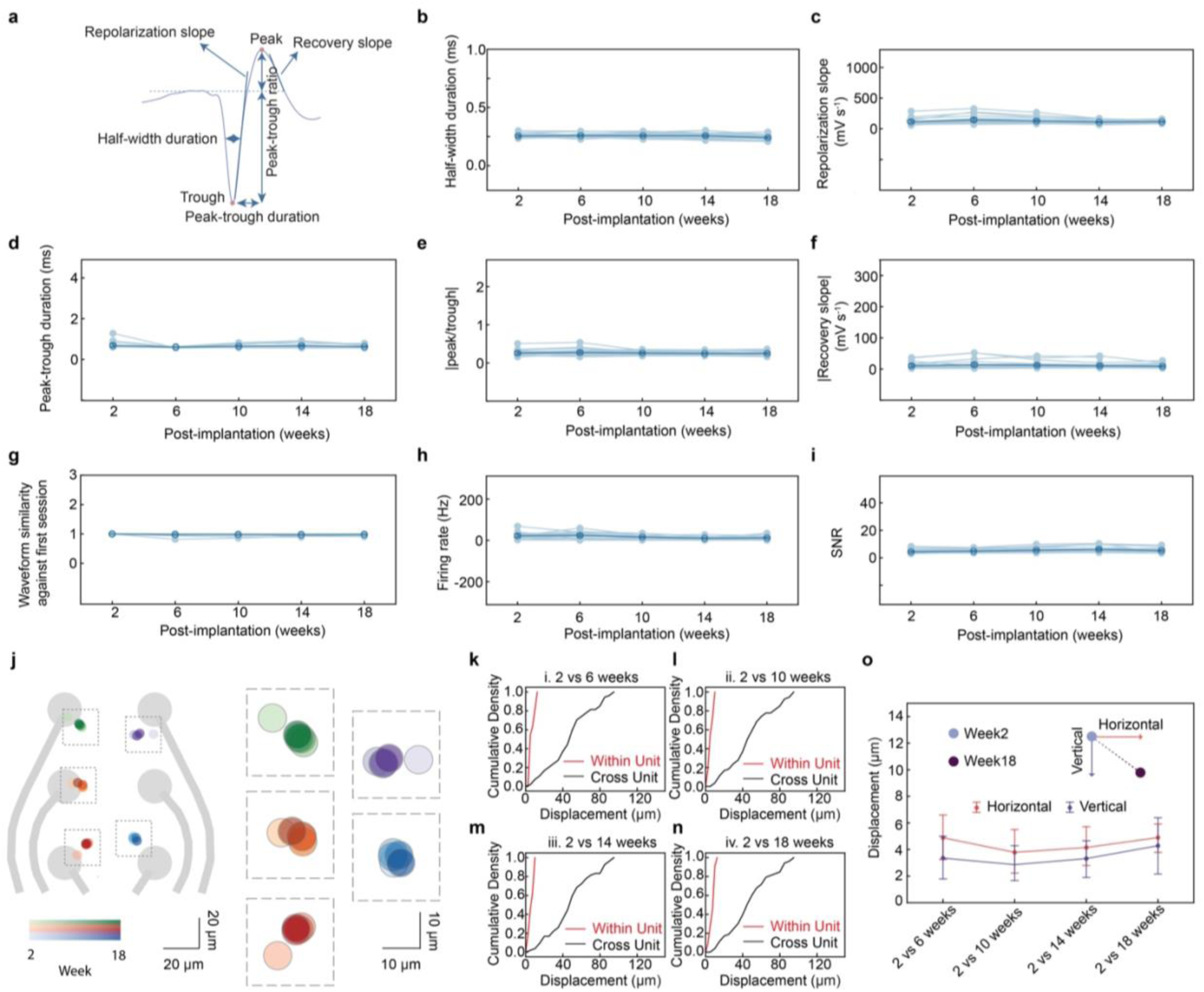
Long-term tracking of the neural activity from the same cells by FlexiSoft electrodes. a-f, Features extracted from the single-unit waveform (a) as a function of recording weeks, including half-width duration (b), repolarization slope (c), peak-trough duration (d), absolute peak-trough ratio (e), and absolute recovery slope (f). g-i, Waveform similarity (g), firing rate (h), and SNR (i) for different units as a function of recording time. One-way ANOVA followed by Tukey correction showing non-significant changes, data represented as mean ± s.d., *n* = 22 neurons, *p* > 0.05. j, Single-neuron waveform centroids throughout 18-week recording from one shank of a representative mouse. The centroid for each single neuron is shown for week 2, 6, 10, 14, and 18 with gradient colors. Gray circles indicate electrode position in one shank. k-n, Cumulative distribution of within-unit centroid displacement (red) between week 2 and weeks 6, 10, 14, and 18, compared to across-unit displacement within a day (black). For all comparisons of within-unit displacement, *P* < 0.001 (two-sided Wilcoxon rank-sum test). o, Statistical summary of vertical and horizontal displacement of the same neuron across the entire recording period. Data represented as mean ± SD, *n* = 22 neurons from 1 representative mouse.

**Extended Data Fig. 6.**
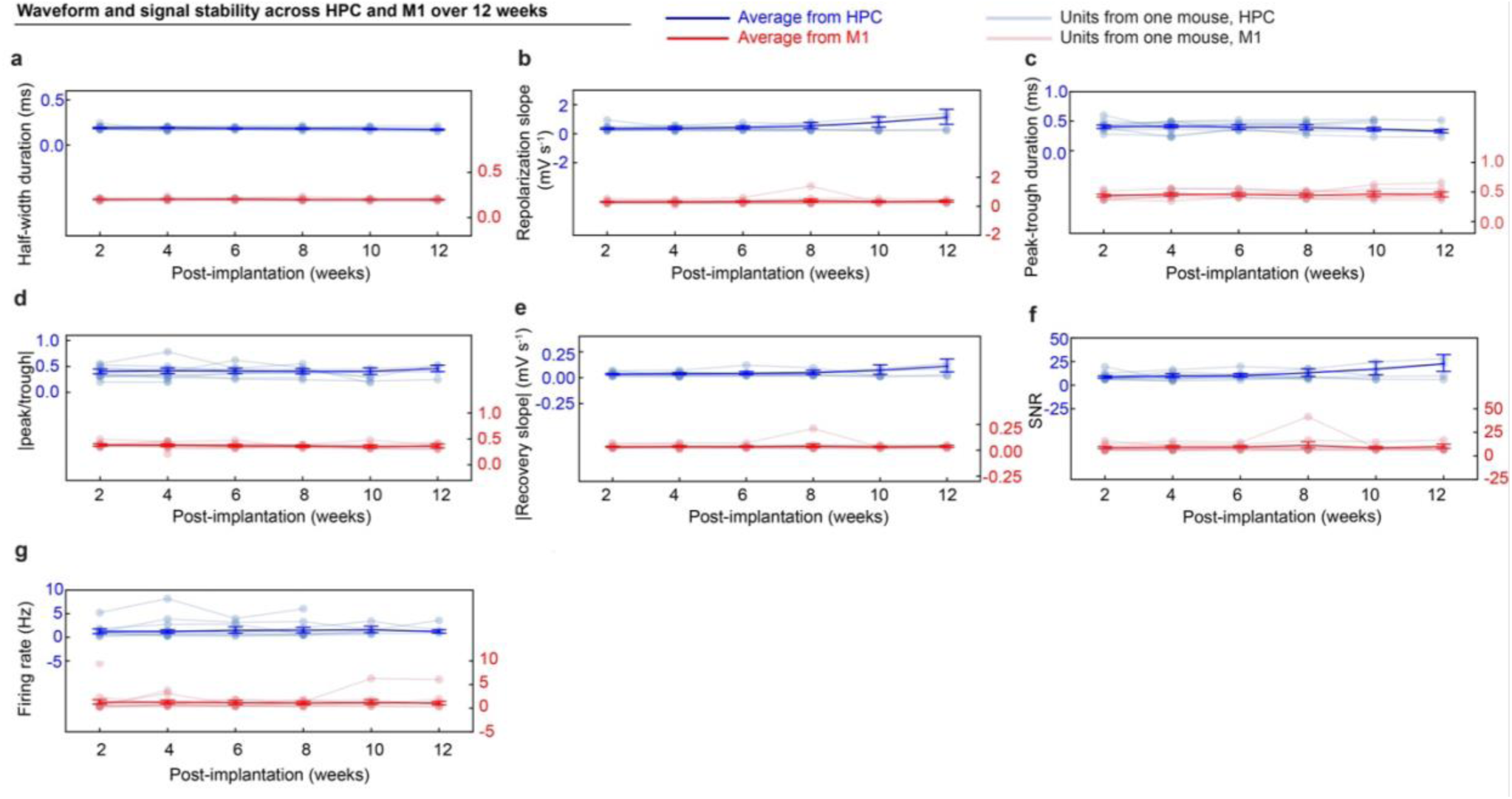
Long-term tracking of neural activity from the same cells across HPC and M1. a-e, Analysis of waveform features—half-width duration (a), repolarization slope (b), peak-trough duration (c), absolute peak-trough ratio (d), and recovery slope (e) — across HPC and M1 units over 12 weeks. f-g, Analysis of SNR (f) and firing rate (g) across 12 weeks post implantation. In total, 156 units from HPC and 203 units from the M1 region from 13 mice were analyzed. One-way ANOVA followed by Tukey correction were conducted for all features. Changes were considered non-significant if *p* > 0.05. In the HPC, all waveform features were stable, with 95.8% of units also showing no changes in firing rate and SNR. In M1, all waveform features except peak-trough duration were stable across units; 97.3% showed non-significant changes in peak-trough duration, firing rate, and SNR.

**Extended Data Fig. 7.**
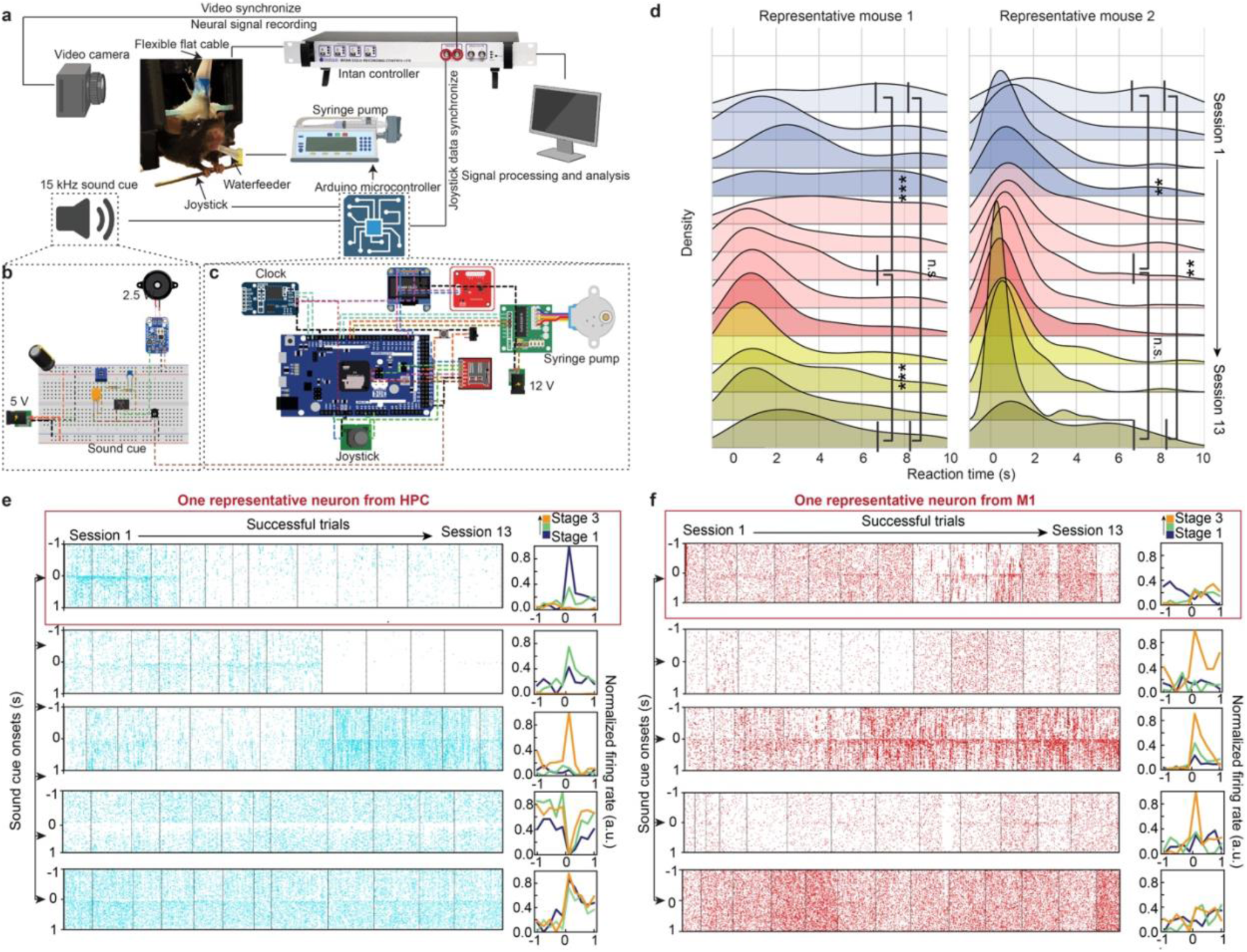
Long-term tracking neural activity from the same cells during a motor learning task by FlexiSoft probes. a-c, Schematic of the integrated neural signal recording and behavioral tracking system for joystick-based learning tasks in mice. The setup includes a neural recording system (Intan), joystick interface, syringe pump for water reward delivery, sound cue generator, and video camera, all synchronized by an Arduino-based controller. d, Density plots of reaction times between sound cue onset and the time when the joystick crossed the threshold, indicating trial success, across sessions. Unpaired, two-tailed t-test, *n* = 100 trials, * *p* < 0.05, ** *p* < 0.01, *** *p* < 0.001. e-f, Spike raster plots during successful trials across all 13 training sessions for 5 representative neurons from the HPC (e) and M1 (f). Insets show the normalized firing rates from stage 1 (sessions 1-4), stage 2 (sessions 5-9), and stage 3 (sessions 10-13) training sessions for the corresponding neurons.

**Extended Data Fig. 8.**
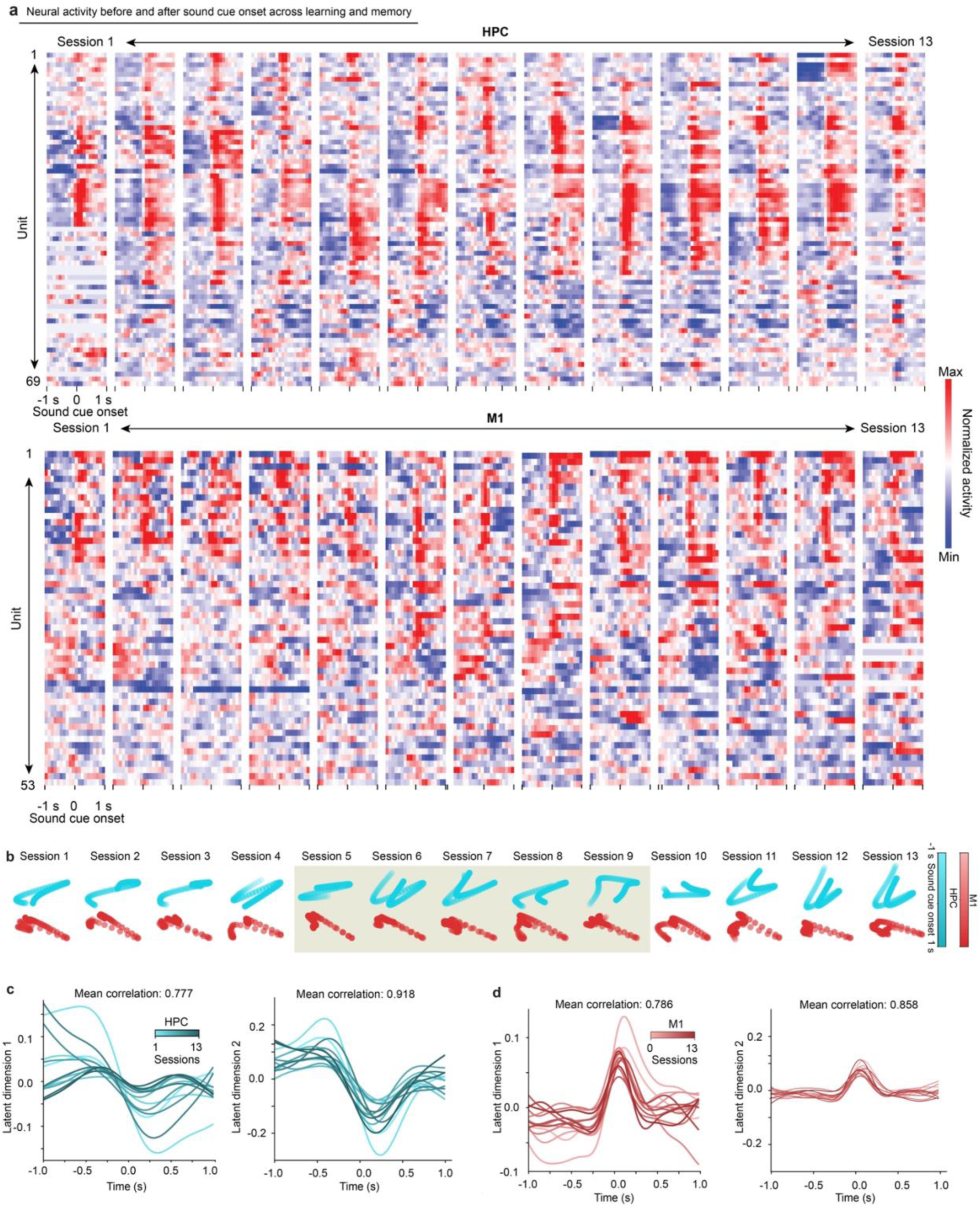
Population analysis of neural activity from the same cells during motor learning. a, Normalized neural activity heatmaps for all the neurons from 5 representative mice in the HPC (upper) and M1 (lower) regions over 13 training sessions. b, The latent dynamics from one representative mouse averaged across all trial repeats (100 trials per session) in each recording session over the entire training period from HPC (blue) and M1 region (red). c-d, Latent dynamics projected onto the first and second latent dimensions for the HPC (c) and M1 (d) from one representative mouse.

**Extended Data Fig. 9.**
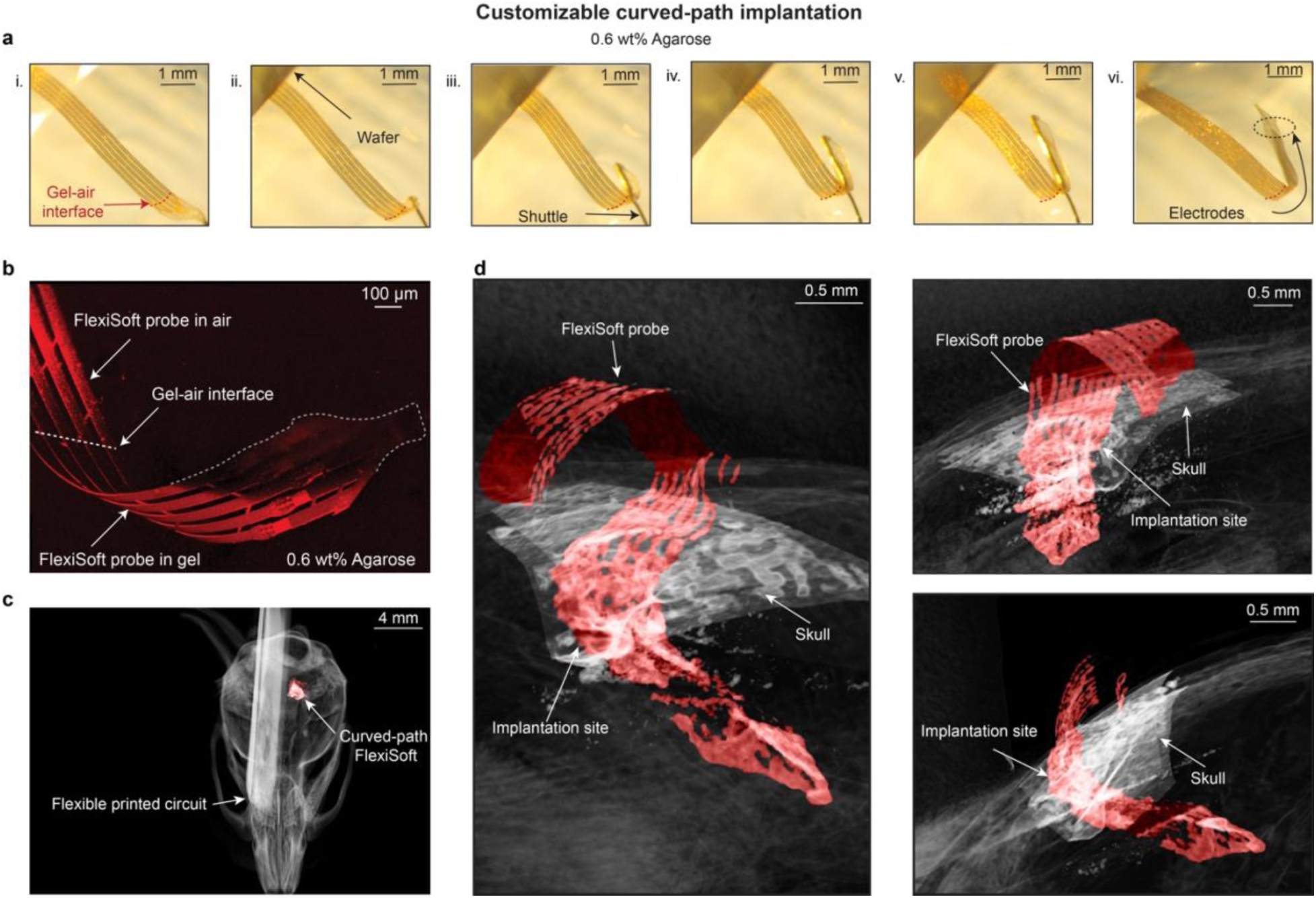
Curved-path implantation. a, BF images showing the stepwise curved- path implantation of a FlexiSoft probe in a 0.6 wt% agarose gel. b, 3D reconstructed confocal fluorescence image of the FlexiSoft probe in the agarose gel following the curved-path implantation. c-d, Micro-CT images showing dorsal-to-ventral (c) and lateral views (d) of a representative FlexiSoft probe implanted into a mouse brain following curved-path implantation.

**Extended Data Fig. 10.**
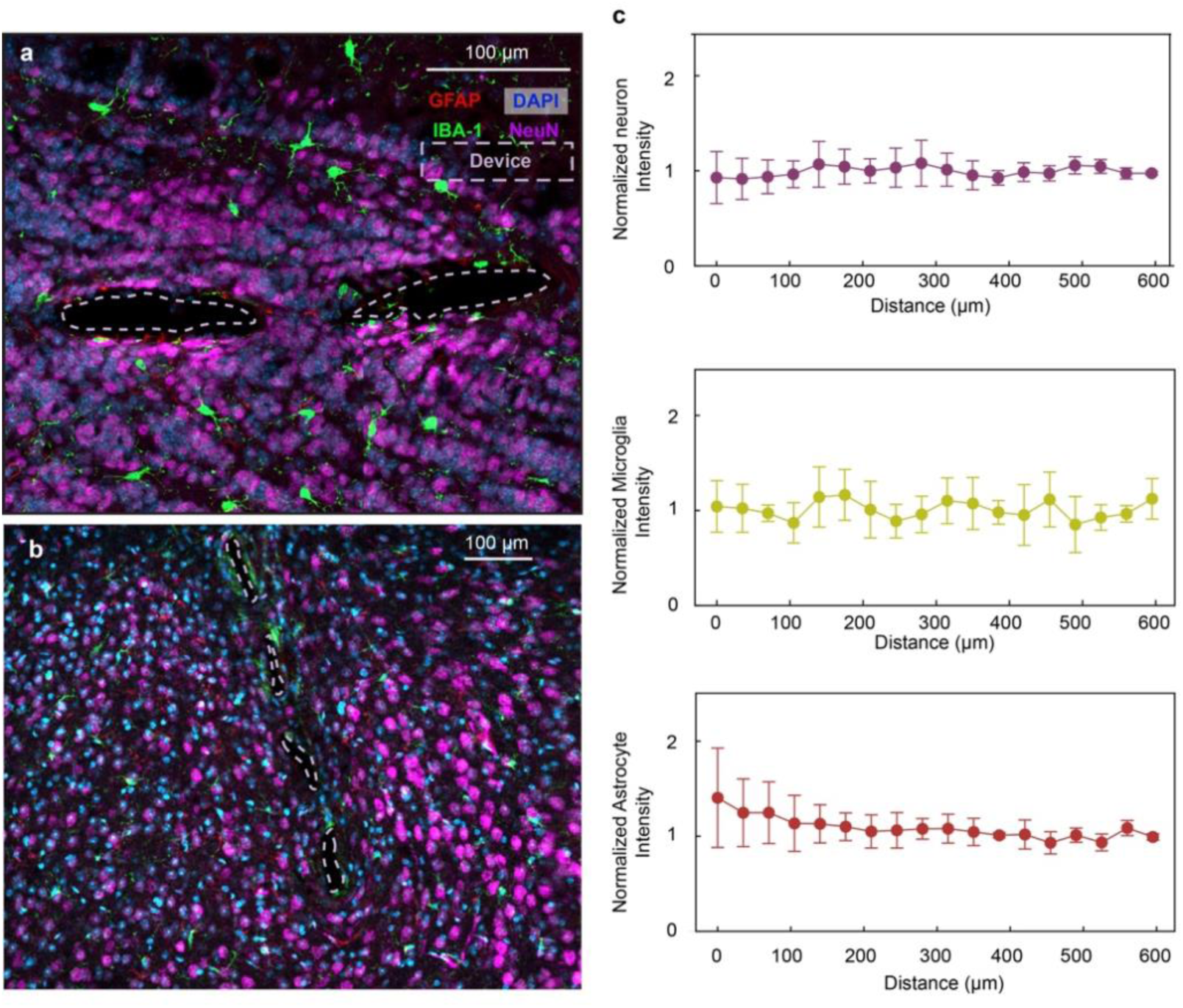
Immunostaining characterization. a-b, Representative fluorescence images showing the immunostaining results of brain slices from olfactory bulb (a) and NAc (b) regions after 6 months post-implantation showing the cross-section of FlexiSoft probes. c, Normalized average fluorescence intensity of NeuN, IBA-1, and GFAP signals plotted as a function of distance from the probe–tissue interfaces (*n* = 6 brain slices from 2 mice).

## Supplementary information

The supplementary information document contains **Supplementary Discussion 1-4** and **Supplementary Figure 1-2**.

### Supplementary Discussion 1: Energy release rate of a homogeneous neural probe

We first calculate the stress intensity factor *KI* (Supplementary Fig. 1a), considering a rectangular plate with length 2*l*, width *b* and thickness *h*, containing an edge crack of length *a* that is unconstrained. For the plate dimensions such that the length-to-width ratio *l*/*b* ≥ 0.5 and the crack length-to-width ratio *a*/*b* ≤ 0.6, the stress intensity factor *KI* at the crack tip under a uniaxial stress

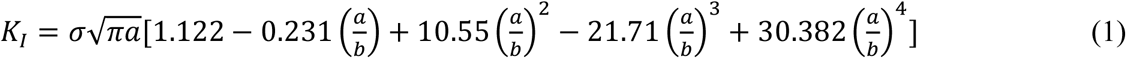

Second, we calculate energy release rate *G*, which is directly related to the Mode-I stress intensity factor *KI*. For a crack propagating under a Mode-I (opening mode) condition in a linearly elastic material, the relationship between *G* and *KI* is established in the context of plane stress. The *G* for Mode-I is given by:

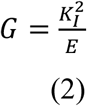

where *E* is the Young’s modulus of the material. This formulation is appropriate for the current analysis, as the crack is modeled under plane stress conditions.

Taking equation (1) into equation (2) we can get the solution for our energy release rate as:

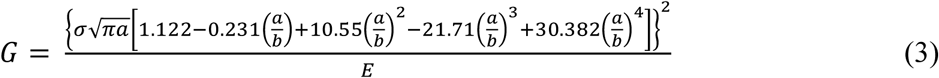

The simulation here and the solution from other literature^1^ for homogeneous neural probe align well with each other (Supplementary Fig. 1b). For theoretical calculations of energy release rate for multilayer structure, please refer to other literatures^2–4^.

### Supplementary Discussion 2: Flexural rigidity of a homogeneous neural probe

The flexural rigidity of a single-layer neural probe along its longitudinal axis can be approximated using the Euler-Bernoulli beam model, assuming the lateral dimensions are significantly larger than the thickness. For a homogeneous beam, the flexural rigidity *D* is given by the following equation:

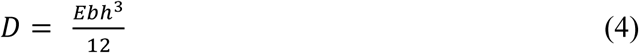

where *E* is the elastic modulus of the material, *b* is the width of the beam and *h* is the thickness of the beam.

### Supplementary Discussion 3: Flexural rigidity of a two-layer heterogeneous neural probe

To analyze the flexural rigidity of the two-layer neural probe, we first determine the location of the neutral axis. At the neutral axis, the sum of forces in the x-direction must be zero, and the sum of moments around the z-axis must also be zero (Supplementary Fig. 1c):

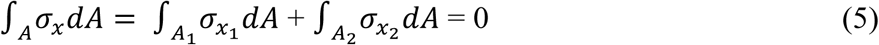

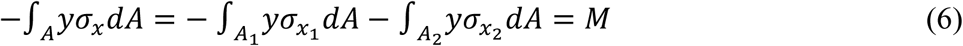

where *A*, *A1* and *A2* represent the cross-sectional areas of the entire structure, H-SEBS, and SU-8, respectively. Similarly, *σx*, *σx1*, and *σx2* denote the normal stress in the x direction for the entire structure, H-SEBS, and SU-8, respectively. The variable *y* represents the distance from a given point to the neutral axis, and *M* represents the moment with respect to that point.

Both materials composing the beam obey Hooke’s law, with elastic modulus *E1* for H-SEBS and *E2* for SU-8. The normal stress varies linearly with the distance *y* from the neutral axis:

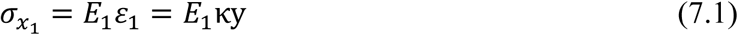

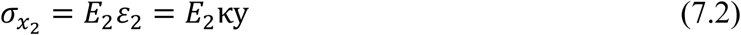

where *k* is the curvature of the beam, *ε1* and *ε2* denote the normal strain in the x direction for the H-SEBS and SU-8, respectively. We also introduce notation here:

Substituting (7) and (8) into (5) to obtain:

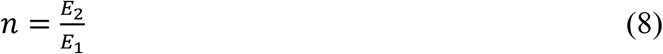

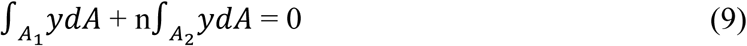

Using the top of H-SEBS film as a reference, from (9) we can get the location of the neutral axis:

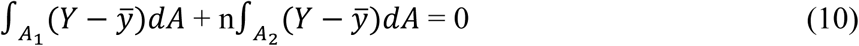

where *Y* is the distance from a point in the structure to the neutral axis. *y* represents the location of the neutral axis of the entire structure relative to the top of the H-SEBS film. It is the weighted average of the centroidal positions of the H-SEBS and SU-8 layers. According to equation (10) we have:

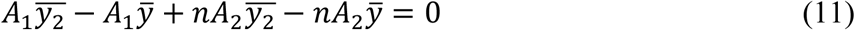

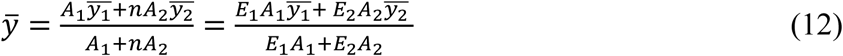

where *y*_1_ and *y*_2_ refers to the centroidal distance from the top of the H-SEBS film and SU-8 film to the neutral axis of the top of each layer respectively. Substituting the width b for both H-SEBS and SU-8 and thickness *h1* and *h2* for H-SEBS and SU-8, respectively, we can further change equation (12) into:

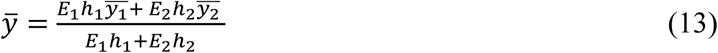

Then, total flexural rigidity can be written as:

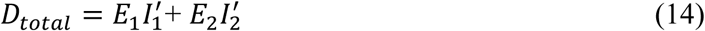

Where *I1* and *I2* represent the original moment of inertia for the H-SEBS and SU-8 layers, respectively, calculated based on their geometry about their own centroidal axes. They represent the inherent resistance of each material layer to bending around their own centroidal axis. When we shift coordinates to the neutral axis, according to the parallel axis theorem, the moment of inertia shifts, giving us *I1*′ and *I2*′, which include the effect of the offset between the neutral axis and each layer’s own centroidal axis. These shifted moment of inertia values are calculated as:

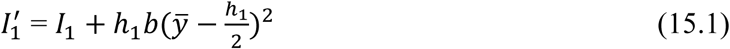

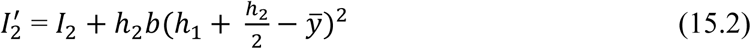

Taken equation (15) into equation (14) we can get:

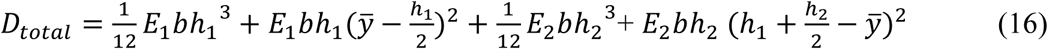

### Supplementary Discussion 4: Set-up of learning tasks

The experimental setup includes a joystick and a syringe pump actuator (water feeder), both controlled by an Arduino microcontroller, as well as a sound cue device used for training the mice, a camera for recording training videos, and a neural signal acquisition system (Intan Technologies) for recording neural signals (Extended Data Fig. 8a). To synchronize the data from joystick displacement, videos, and neural signals, binary trigger signals are sent from both the camera and Arduino microcontroller to the Intan system, ensuring precise alignment for post-experiment analysis. The sound cue device operates using a 555 timer integrated circuit (IC), which generates a constant pulse-width modulation (PWM) signal. This signal is amplified to produce a consistent auditory cue (Extended Data Fig. 8b). The syringe pump is driven by a stepper motor that controls the movement of a lead screw, which in turn operates the syringe to deliver water rewards (Extended Data Fig. 8c). The joystick functions as an analog system, and its position data is processed into angle and distance information for analysis. The entire system runs on a state machine program that optimizes the operation of all peripherals, ensuring efficient performance without blocking delays. Additionally, the setup includes a real-time clock (RTC) for accurate time synchronization of data storage, a near-field communication (NFC) card for recording the subject’s ID, and an organic light-emitting diode (OLED) display to provide real-time updates on the experiment’s status. User interaction is facilitated through control switches, allowing adjustments during the experiment.

---

**Supplementary Information Fig. 1.**
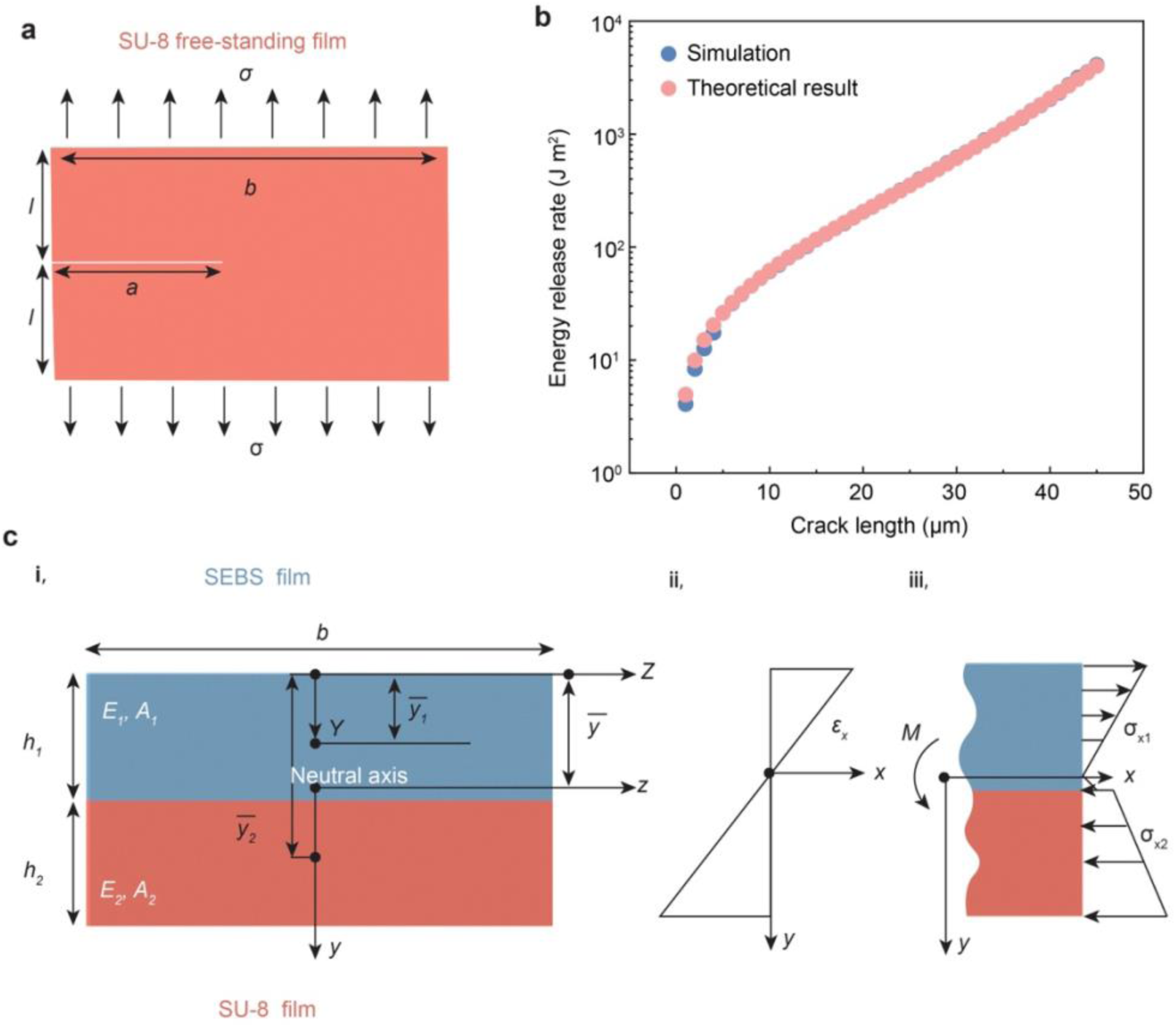
Simulation and theoretical calculation for energy release rate and flexural rigidity. a, A free-standing SU-8 film with an edge crack of length *a* under uniaxial tensile stress *σ*. This configuration is used to model the energy release rate during crack propagation. b, Simulated and Theoretical result of energy release rate as a function of crack length for single-layer SU-8 structure. c, An SU-8/H-SEBS multilayer composite model for flexural rigidity analysis: i, Cross-sectional view of the H-SEBS (blue) and SU-8 (red) layers with respective thicknesses *h1* and *h2*, elastic modulus *E1* and *E2*, and areas *A1* and *A2*. The neutral axis is located between the layers, with centroidal distances *y*_1_ and *y*_2_from the top of each layer. ii, Strain distribution across the cross section, showing linear variation of strain *εx* with respect to the distance from the neutral axis. iii, Stress distribution under applied moment *M*, showing tensile stress *σx1* in the H-SEBS layer and compressive stress *σx2* in the SU-8 layer.

**Supplementary Information Fig.2.**
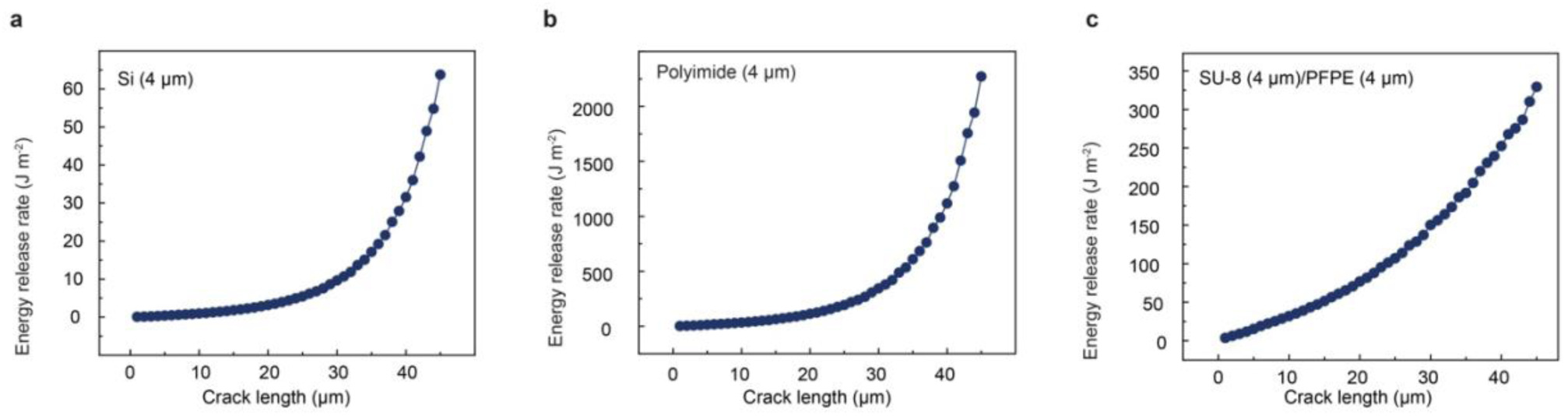
Simulation results depicting the energy release rate for neuralprobe with different materials: a, 4-μm-thick silicon rigid layer; b, 4-μm-thick Polyimide (PI) flexible layer; c, 4-μm-thick SU-8/4-μm-thick PFPE hybrid layer.

## References

1 Kim, J. H., Daie, K. & Li, N. A combinatorial neural code for long-term motor memory. Nature 637, 663–672 (2025).

2 Grewe, B. F. et al. Neural ensemble dynamics underlying a long-term associative memory. Nature 543, 670–675 (2017).

3 Williamson, M. R. et al. Learning-associated astrocyte ensembles regulate memory recall. Nature 637, 478–486 (2025).

4 Gallego, J. A., Perich, M. G., Chowdhury, R. H., Solla, S. A. & Miller, L. E. Long-term stability of cortical population dynamics underlying consistent behavior. Nat. Neurosci. 23, 260–270 (2020).

5 Cho, Y., Park, S., Lee, J. & Yu, K. J. Emerging materials and technologies with applications in flexible neural implants: A comprehensive review of current issues with neural devices. Adv. Mater. 33, e2005786 (2021).

6 Eimon, P. M. et al. Brain activity patterns in high-throughput electrophysiology screen predict both drug efficacies and side effects. Nat. Commun. 9, 219 (2018).

7 Jun, J. J. et al. Fully integrated silicon probes for high-density recording of neural activity. Nature 551, 232–236 (2017).

8 Steinmetz, N. A. et al. Neuropixels 2.0: A miniaturized high-density probe for stable, long- term brain recordings. Science 372, eabf4588 (2021).

9. 9 Ye, Z., et al. Ultra-high density electrodes improve detection, yield, and cell type identification in neuronal recordings. bioRxiv 10.1101/2023.08.23.554527 (2024).

10 Buccino, A. P., Garcia, S. & Yger, P. Spike sorting: New trends and challenges of the era of high-density probes. *Prog*. Biomed. Eng. 4, 022005 (2022).

11 Tang, X., Shen, H., Zhao, S. Y., Li, N. & Liu, J. Flexible brain-computer interfaces. Nat. Electron. 6, 109–118 (2023).

12 Remy, A., Lin, X. Y. & Liu, J. Materials for flexible and soft brain-computer interfaces, a review. MRS Commun. 14, 827–834 (2024).

13 Barz, F., Ruther, P., Takeuchi, S. & Paul, O. Mechanically adaptive silicon-based neural probes for chronic high-resolution neural recording. Procedia Eng. 120, 952–955 (2015).

14 Otte, E., Cziumplik, V., Ruther, P. & Paul, O. Customized thinning of silicon-based neural probes down to 2 microm. Annu. Int. Conf. IEEE Eng. Med. Biol. Soc. 2020, 3388–3392 (2020).

15 HajjHassan, M., Chodavarapu, V. & Musallam, S. NeuroMEMS: Neural probe microtechnologies. Sensors 8, 6704–6726 (2008).

16 Zhao, S. Y. et al. Tracking neural activity from the same cells during the entire adult life of mice. Nat. Neurosci. 26, 1129–1129 (2023).

17 Le Floch, P. et al. 3D spatiotemporally scalable in vivo neural probes based on fluorinated elastomers. Nat. Nanotechnol. 19, 319–329 (2024).

18 Oh, S. et al. A stealthy neural recorder for the study of behaviour in primates. *Nat*. Biomed. Eng. 10.1038/s41551-024-01280-w (2024).

19 Shin, H. et al. Multifunctional multi-shank neural probe for investigating and modulating long-range neural circuits *in vivo*. Nat. Commun. 10, 3777 (2019).

20 Jeon, M. et al. Partially flexible MEMS neural probe composed of polyimide and sucrose gel for reducing brain damage during and after implantation. J. Micromech. Microeng. 24, 025010 (2014).

21 Zhang, S., Wang, C. J., Linghu, C. H., Wang, S. H. & Song, J. Z. Mechanics strategies for implantation of flexible neural probes. J. Appl. Mech. 88, 010801 (2021).

22 You, X. L. et al. Progress in mechanical modeling of implantable flexible neural probes. Comp. Model. Eng. 140, 1205–1231 (2024).

23 Douville, N. J., Li, Z. Y., Takayama, S. & Thouless, M. D. Fracture of metal coated elastomers. Soft Matter 7, 6493–6500 (2011).

24 Vlassak, J. J. Channel cracking in thin films on substrates of finite thickness. Int. J. Fracture 119, 299–323 (2003).

25 Huang, R., Prévost, J. H., Huang, Z. Y. & Suo, Z. Channel-cracking of thin films with the extended finite element method. Eng. Fract. Mech. 70, 2513–2526 (2003).

26 Beuth, J. L. Cracking of thin bonded films in residual tension. Int. J. Solids Struct. 29, 1657–1675 (1992).

27 Cai, Y. J. et al. Enhancing the fracture resistance of hydrogels by regulating the energy release rate via bilayer designs: Theory and experiments. J. Mech. Phys. Solids 170, 105125 (2023).

28 Tan, E. K. W. et al. Nanofabrication of conductive metallic structures on elastomeric materials. Sci. Rep. 8, 6607 (2018).

29 Criscuolo, V. et al. Double-framed thin elastomer devices. ACS Appl. Mater. Inter. 12, 55255–55261 (2020).

30 Chen, R., Canales, A. & Anikeeva, P. Neural recording and modulation technologies. Nat. Rev. Mater. 2, 16093 (2017).

31 Han, J. et al. Impact of impedance levels on recording quality in flexible neural probes. Sensors 24, 2300 (2024).

32 Chung, J. E. et al. A fully automated approach to spike sorting. Neuron 95, 1381–1394 (2017).

33. 33 McInnes, L., Healy, J. & Melville, J. Umap: Uniform manifold approximation and projection for dimension reduction. arXiv 10.48550/arXiv.1802.03426 (2018).

34 Rosenkranz, K., Kacar, A. & Rothwell, J. C. Differential modulation of motor cortical plasticity and excitability in early and late phases of human motor learning. J. Neurosci. 27, 12058–12066 (2007).

35 van Beest, E.H., Bimbard, C., Fabre, J.M.J. et al. Tracking neurons across days with high- density probes. Nat. Methods 22, 778–787 (2025).

36 Schoonover, C. E., Ohashi, S. N., Axel, R. & Fink, A. J. P. Representational drift in primary olfactory cortex. Nature 594, 541–546 (2021).

37 Kim, J., Joshi, A., Frank, L. & Ganguly, K. Cortical-hippocampal coupling during manifold exploration in motor cortex. Nature 613, 103–110 (2023).

38 Lee, C., Harkin, E. F., Yin, X. M., Naud, R. & Chen, S. Cell-type-specific responses to associative learning in the primary motor cortex. Elife 11, e72549 (2022).

39 Hwang, E. J., Dahlen, J. E., Mukundan, M. & Komiyama, T. Disengagement of motor cortex during long-term learning tracks the performance level of learned movements. J. Neurosci. 41, 7029–7047 (2021).

40 Bolin, T. et al. Automated home cage training of mice in a hold-still center-out reach task. J. Neurophysiol. 121, 500–512 (2019).

41 Hwang, E. J. et al. Disengagement of motor cortex from movement control during long- term learning. Sci. Adv. 5 (2019).

42 Varani, A. P. et al. Multiple functions of cerebello-thalamic neurons in learning and offline consolidation of a motor skill in mice. *bioRxiv*, 10.1101/2024.09.11.612475 (2024).

43 Laeremans, A. et al. Distinct and simultaneously active plasticity mechanisms in mouse hippocampus during different phases of Morris water maze training. Brain Struct. Funct. 220, 1273–1290 (2015).

44 Dolfen, N., Reverberi, S., Op de Beeck, H., King, B. R. & Albouy, G. The hippocampus represents information about movements in their temporal position in a learned motor sequence. J. Neurosci. 44, e0584242024 (2024).

45 Yu, B. M. et al. Gaussian-process factor analysis for low-dimensional single-trial analysis of neural population activity. J. Neurophysiol. 102, 2008–2008 (2009).

46 Buccino, A. P. et al. SpikeInterface, a unified framework for spike sorting. Elife 9, e61834 (2020).

47 Zhao, Z. T. et al. Ultraflexible electrode arrays for months-long high-density electrophysiological mapping of thousands of neurons in rodents. *Nat*. Biomed. Eng. 7, 520–532 (2023).

48 Gao, L. et al. Free-standing nanofilm electroe arrays for long-term stable neural interfacings. Adv. Mater. 34, 2107343 (2022).

49 Liu, Y. et al. A high-density 1,024-channel probe for brain-wide recordings in non-human primates. Nat. Neurosci. 27, 1620–1631 (2024).

50 Wang, X. C. et al. A parylene neural probe array for multi-region deep brain recordings. J. Microelectromech. S. 29, 499–513 (2020).

51 Khatib, M. et al. Spiral NeuroString: High-density soft bioelectronic fibers for multimodal sensing and stimulation. *bioRxiv* 10.1101/2023.10.02.560482 (2023).

## Additional References

1 Liu, M., Gan, Y., Hanaor, D. A., Liu, B. & Chen, C. An improved semi-analytical solution for stress at round-tip notches. Eng. Fract. Mech. 149, 134–143 (2015).

2 Beuth, J. L. Cracking of thin bonded films in residual tension. Int. J. Solids Struct. 29, 1657–1675 (1992).

3 Huang, R., Prévost, J. H., Huang, Z. Y. & Suo, Z. Channel-cracking of thin films with the extended finite element method. Eng. Fract. Mech. 70, 2513–2526 (2003).

4 Vlassak, J. J. Channel cracking in thin films on substrates of finite thickness. Int. J. Fracture 119, 299–323 (2003).

